# Modelling Porcine NAFLD by Deletion of Leptin and defining the role of AMPK in hepatic fibrosis

**DOI:** 10.1101/2021.06.07.447327

**Authors:** Tan Tan, Zhiyuan Song, Runming Wang, Shuheng Jiang, Zuoxiang Liang, Qi Wang, Xiaoxiang Hu, Ning Li, Yiming Xing

**Author notes:** Correspondence (Y. Xing). These authors contribute equally to this work (Tan Tan, Zhiyuan Song).

## Abstract

Liver fibrosis occurs during chronic liver disease. Advanced liver fibrosis results in cirrhosis, liver failure and often requires liver transplantation. However, due to the lack of human models, mechanisms underlining the pathogenesis of liver fibrosis remain unclear. Recent studies implicated a central role of deranged lipid metabolism in its pathogenesis. In this study, we generated LEPTIN-deficient (*LEPTIN^-/-^*) pigs using zinc finger nuclease technology to investigate the mechanisms of liver fibrosis associated with obesity. The *LEPTIN^-/-^* pigs showed increased body fat and significant insulin resistance by 12 months of age. To resemble non-alcoholic fatty liver disease (NAFLD) patients, *LEPTIN^-/-^* pig developed the phenotypic features of fatty liver, non-alcoholic steatohepatitis (NASH) and hepatic fibrosis with age. Meanwhile, LEPTIN absence reduced phosphorylation of JAK2-STAT3 and AMPK. The alteration of JAK2-STAT3 enhanced fatty acid β-oxidation, whereas inactivation of AMPK led to mitochondrial autophagy, and both contributed to increased oxidative stress in hepatocytes. Although Leptin deletion in the rat liver altered JAK2-STAT3 phosphorylation, it activated the AMPK pathway and prevented liver fibrogenesis in contrast with the *LEPTIN^-/-^* pig. To our knowledge, the *LEPTIN^-/-^* pig provides the first model recapitulating the full pathogenesis of NAFLD and its progression toward liver fibrosis. The activity of AMPK signaling pathway suggests a potential target for development of new strategies for the diagnosis and treatment of NAFLD.

## Introduction

Liver fibrosis is a common result of chronic damage to the liver caused by the accumulation of extracellular matrix proteins. A number of liver diseases as well as side effects of some drugs lead to liver fibrosis. Among these causes, non-alcoholic steatohepatitis (NASH) has been recognized as a major etiology (1). It is considered part of the spectrum of non-alcoholic fatty liver disease (NAFLD) (2). Excessive adipose tissue in NAFLD patients leads to an inflammatory state targeting liver parenchyma with steatosis and fibrosis observed in a significant portion of cases. Hepatic fibrogenesis can progress to cirrhosis which enhances the risk of hepatocellular carcinoma. Alarmingly, few effective pharmacotherapeutic approaches are currently available to block or attenuate development and progression of NAFLD. Liver transplantation is the only efficient treatment option in patients with decompensated cirrhosis (3). Nowadays, the fibrosis stage is viewed as the most important predictor of mortality in NAFLD. However, it is not a unidirectional progressive process, ultimately leading to liver cirrhosis and organ failure, but is in principle reversible. Accordingly, better understanding of mechanisms underlying fibrogenesis are crucial for the development of new strategies for the prevention and treatment of NAFLD and other related liver disease.

Although the pathogenesis of liver fibrosis has not been fully discovered, the primary role is played by the deposition of triglycerides in liver cells and the formation of lipid droplets (4). An accumulation of fatty liver leads to insulin resistance in adipose tissue with increased pro-inflammatory cytokines, which initiate the necrosis and apoptosis of liver cells (5). Elevated free fatty acid (FFA) aggravates the oxidative damage of liver cells, and also leads to insulin resistance. Insulin resistance worsens adipocyte function and promotes the transition of NASH to overt liver fibrosis (6). Although the mechanism of fibrosis deposition has been identified, there are almost no therapies currently available that directly prevent or reverse it. While animal models have been indispensable in further studies, current models possess important limitations which largely restricted the understanding of the underlying mechanism driving fibrosis development and discovery of new diagnostics and therapeutics for NAFLD and other liver disease.

Leptin and its receptor play an important role in driving the formation of liver fibrosis. Leptin is an adipocyte-derived hormone that mediates energy homeostasis in various ways, including regulating energy metabolism, promoting oxidation of FFAs and inhibiting fat synthesis (7). A substantial subset of obese patients have relatively low circulating levels of Leptin. Leptin-deficient (*ob/ob*) mice and Leptin receptor-deficient (Zucker) rats are widely used for studying the mechanisms underlying the role of Leptin in hepatic fibrosis. However, these models only represent fatty liver, but not fibrosis in the spontaneous state. Although compounds such as thioacetamide or carbon tetrachloromethane have been shown to behave as potent hepatotoxins which trigger hepatic injury and lead to development of fibrosis, these models cannot fully reflect the unembellished transformation of steatohepatitis to fibrosis in NAFLD patients, either in disease spectrum or etiology (8, 9).

This study aimed to investigate the mechanisms of liver fibrosis and NAFLD caused by Leptin deficiency. Due to the high similarity in physiology and metabolism between humans and pigs, we generated the *LEPTIN^-/-^* pig, which simulates the progression of liver injury, from fatty liver to NASH and hepatic fibrosis, the general physiological alterations and the pathological patterns of NAFLD patients. Compared with *Leptin^-/-^* rats, we discovered that the alteration of JAK2-STAT3 and AMPK signaling pathways mediates β-oxidation and mitochondrial autophagy respectively in *LEPTIN^-/-^* pigs, which in turn enhanced oxidative stress and promoted the development of fatty liver to fibrosis. The *LEPTIN^-/-^* pig model provides a valuable tool to discover the mechanism of the progression of hepatic fibrosis. Meanwhile, the activity of AMPK signaling pathway suggests a potential target to develop new strategy for the diagnosis and treatment of NAFLD.

## Results

### LEPTIN deletion in pigs causes obesity

To generate LEPTIN-knockout pigs, exon 2 of the porcine *LEPTIN* gene was targeted using ZFNs vectors (Figure 1-figure supplement 1A&B). The mutant pig fetal fibroblasts cell clones were screened and transplanted using somatic cell nuclear transfer to generate transgenic pigs. Through DNA sequencing, various mutants were identified in the transgenic pigs (Figure 1-figure supplement 1C&D). As a matter of convenience, mutants were grouped as *LEPTIN^-/-^* and *LEPTIN^+/-^*. The expression of *LEPTIN* were nearly undetectable in *LEPTIN^-/-^* subcutaneous and visceral fat (Figure 1- figure supplement 2A). The LEPTIN protein was not detected in *LEPTIN^-/-^* serum (Figure 1- figure supplement 2B). In order to determine whether the ZFNs resulted in off-target mutations, the genomic regions with the highest levels of homology were analyzed. No off-target mutations were observed at any of the sites profiled using PCR amplification and sequencing (Table S1).

**Figure 1.**
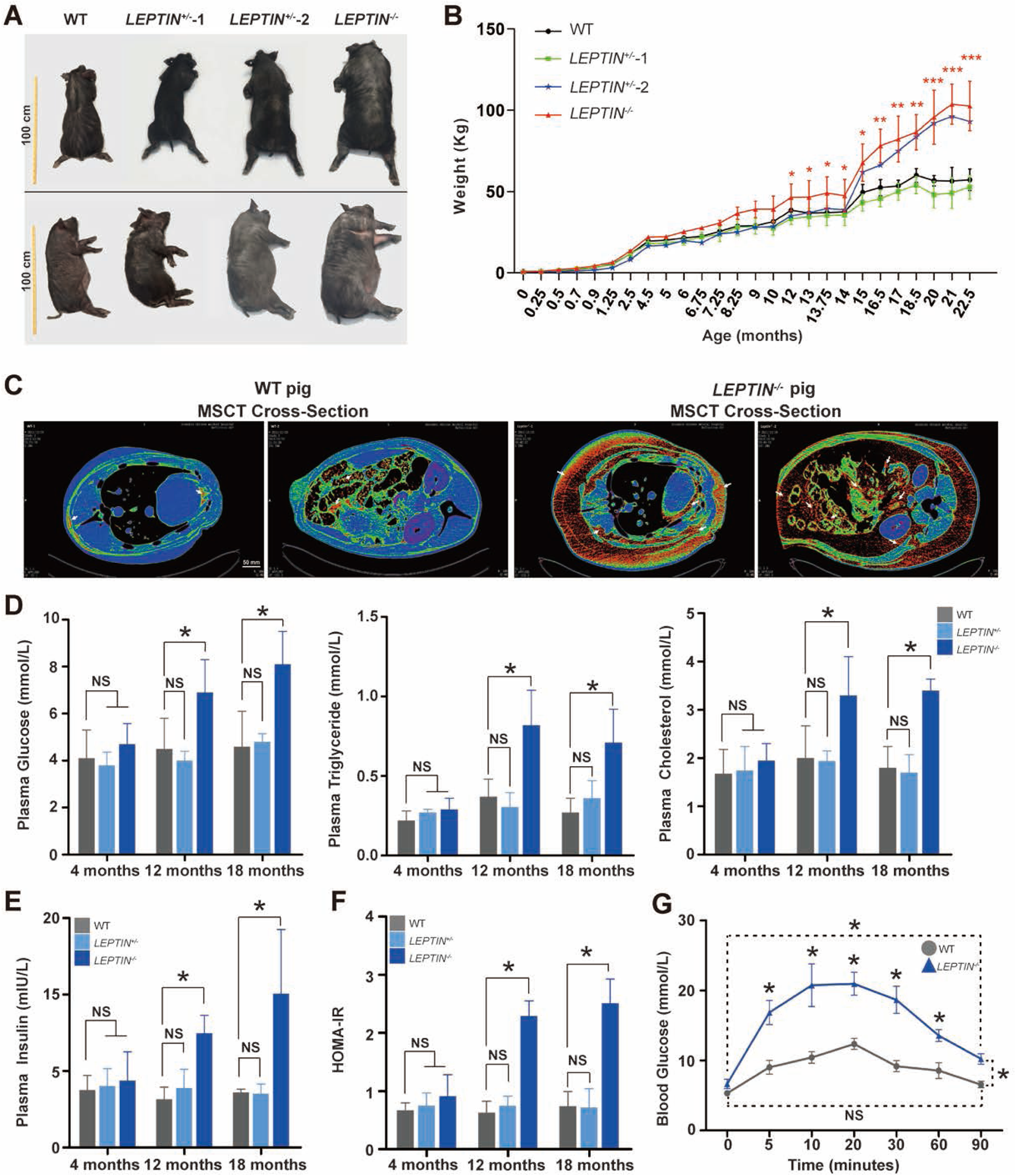
Generation and phenotyping of *LEPTIN*^-/-^ pigs. **A.** Body sizes of *LEPTIN* mutant and WT pigs. Yellow scale bar represents 1meter. **B.** Monthly weight records of pigs. N=3/group. **C.** MSCT scan tomography showing body fat distribution in *LEPTIN* mutant and WT pigs. The red area and white arrows indicated adipose tissue. The scale bar represents 50 mm. **D.** Blood glucose, plasma triglyceride and plasma total cholesterol levels in *LEPTIN* mutant and WT pigs. **E.** Plasma insulin levels in *LEPTIN* mutant and WT pigs. **F.** HOMA-IR results evaluating insulin resistance for pigs at different ages. **G.** IVGTT assay to assess blood glucose regulation. N=3/group. The bars represent the mean±SD; *P<0.05, **P<0.01, and ***P<0.001; NS, non-significant.

Multiple studies have observed that Leptin plays an important role in the development of obesity. The *LEPTIN^-/-^* pigs obtained nearly twice the body weight vs control pigs at 21-months of age (103.75 ± 12.37 kg vs 56.50 ± 8.49 kg) (Figure 1A&B). By MSCT, both subcutaneous and abdominal visceral fat were significantly increased in *LEPTIN^-/-^* compared to control pigs (Figure 1C), and the maximum thickness of subcutaneous fat and the percentage of body fat were more than three times that of control pigs (Figure 1- figure supplement 3A). Using H&E staining *LEPTIN^-/-^* adipocytes also showed increased volumes (Figure 1- figure supplement 3B).

In order to determine whether LEPTIN deficiency altered clinical indicators of liver disease as observed in obese patients, blood was collected from pigs at 4-, 12- and 18-months of age. Compared to control pigs, the concentration of glucose, triglycerides, total cholesterol and low density lipoprotein (LDL) were significantly increased, while high density lipoprotein (HDL) was reduced in *LEPTIN^-/-^* serum at the 12-month and 18-month time points (Figure 1 D & Figure 1- figure supplement 3C). The tendency of these blood tests is similar to that of obese patients which indicate that *LEPTIN^-/-^* pigs demonstrate clinical signs of obesity by 12-months of age.

### LEPTIN deletion in pigs causes type II diabetes

Obesity is one of the main factors contributing to type II diabetes in humans. Consistent with the observed increase in obesity, insulin levels in *LEPTIN^-/-^* pigs were significantly increased at 12- and 18-months of age (Figure 1E). By applying HOMAs to identify diabetic pigs, the insulin and blood glucose parameters indicated that the *LEPTIN^-/-^* pigs were insulin resistance by 12-months of age, with reduced insulin sensitivity and islet β cell function (Figure 1F & Figure 1- figure supplement 4A). Histological analysis demonstrated that the size of *LEPTIN^-/-^* islets and the number of islet β cell were also increased (Figure 1- figure supplement 4B).

Abnormal glucose metabolism is an essential feature of type II diabetes mellitus patients. Thus IVGTT was conducted to determine the pig’s ability to regulate blood glucose. Following the injection of highly concentrated glucose, the glucose concentration in *LEPTIN^-/-^* pigs instantaneously increased to almost 20mmol/L in 10 minutes, which was significantly higher than WT pigs. By 90 minutes post injection, the blood glucose concentration in *LEPTIN^-/-^* pigs was still higher than initial levels (10mmol/L), whereas the WT pigs were nearly fully recovered (Figure 1G). Thus, the lack of LEPTIN negatively affects insulin mediated glucose metabolism, further obstructing glucose regulation and contributing to type II diabetes.

### LEPTIN deletion in pigs results in NAFLD

The incidence rate of non-alcoholic liver injury in obese patients is as high as 90% (12). By the age of 0-6 months, the H&E staining and oil red O staining results showed no visible morphological differences between *LEPTIN^-/-^* and WT livers (normal/total individuals=3/3, 100%) (Figure 2A). However, by the age of 6-12 months, the lipid deposition and hepatocyte steatosis in *LEPTIN^-/-^* pig livers increased (injured/total individuals=3/5, 60%) (Figure 2B). In addition, the triglycerides and the expression of FFA synthesis related genes (*FABP1*, *FASN*, *ELOVL6*, *SCD1* and *PPARA*) were increased significantly in *LEPTIN^-/-^* pigs (Figure 3A&B). H&E and PAS staining of *LEPTIN^-/-^* pig livers at 12-22 months of age revealed features of early stage fatty liver disease; whereas the hepatocytes demonstrated obvious balloon degeneration and vacuolated necrosis consistent with middle and late stages. In addition, a large number of mononuclear cells infiltrated the hepatic lobule portal area and between the hepatic parenchymal cells of hepatic lobules. The PAS assay showed carbohydrate components accumulated in *LEPTIN^-/-^* pig livers (Figure 2C). Real-time PCR results confirmed the expression of *TNFA*, *IL6, NFKB, IL1Β* and *MCP1*, a series of cytokines, were significantly up-regulated in the *LEPTIN^-/-^* pig liver (Figure 3C). IL-1β, TNF-α and IL6 serum concentrations were also significantly increased (damaged individuals/total individuals=2/6, 33%) (Figure 3D).

**Figure 2.**
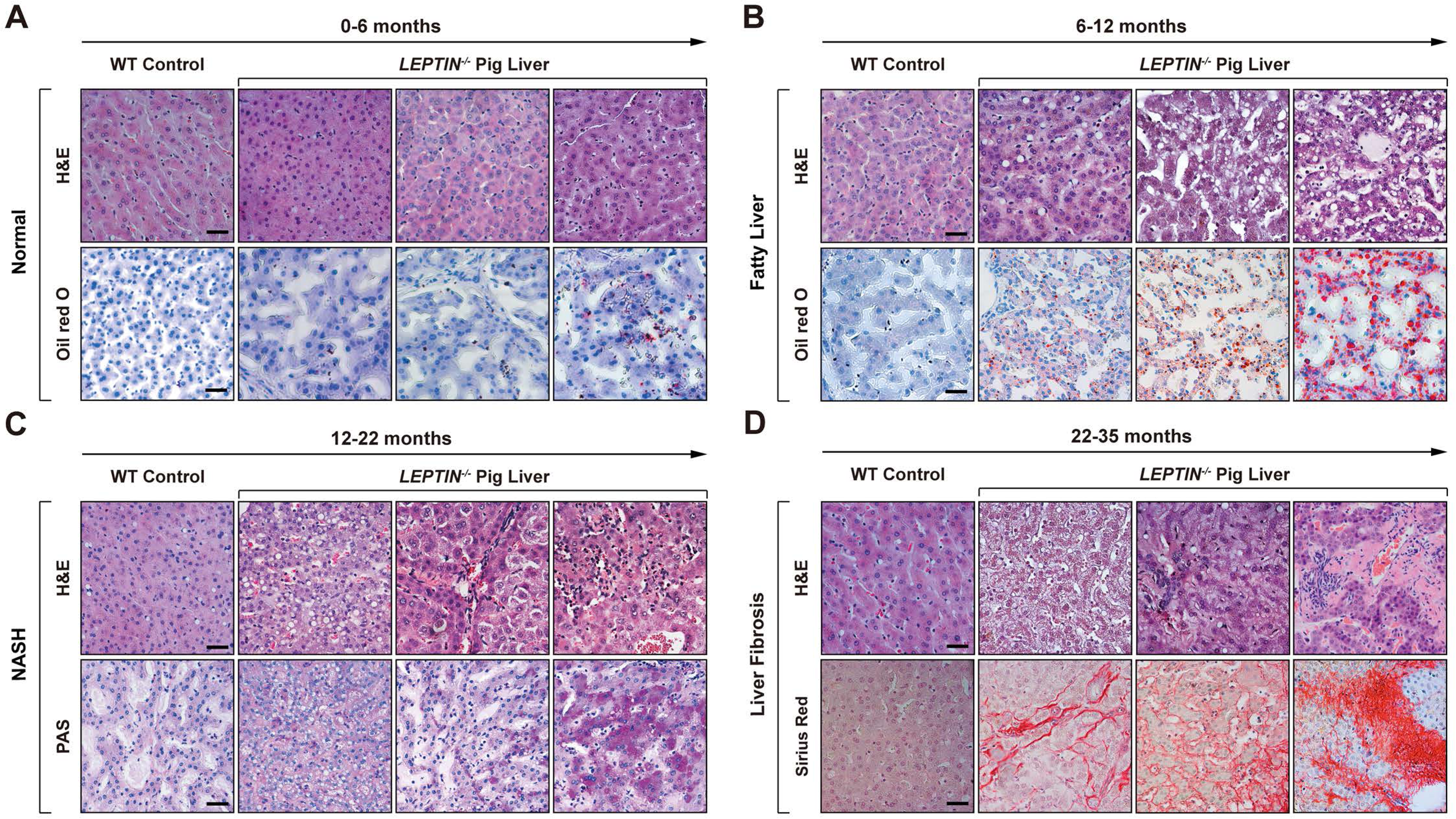
Progression of liver injury in *LEPTIN*^-/-^ pigs. H&E and stage-specific staining demonstrating histological alterations and injury of hepatocytes in *LEPTIN* mutant and WT pigs over time. Oil red O staining for lipid deposition (**A**&**B**), PAS staining for glycogen storage (**C**), and sirius red staining for collagen deposition (**D**). Scale bar represents 50μm.

**Figure 3.**
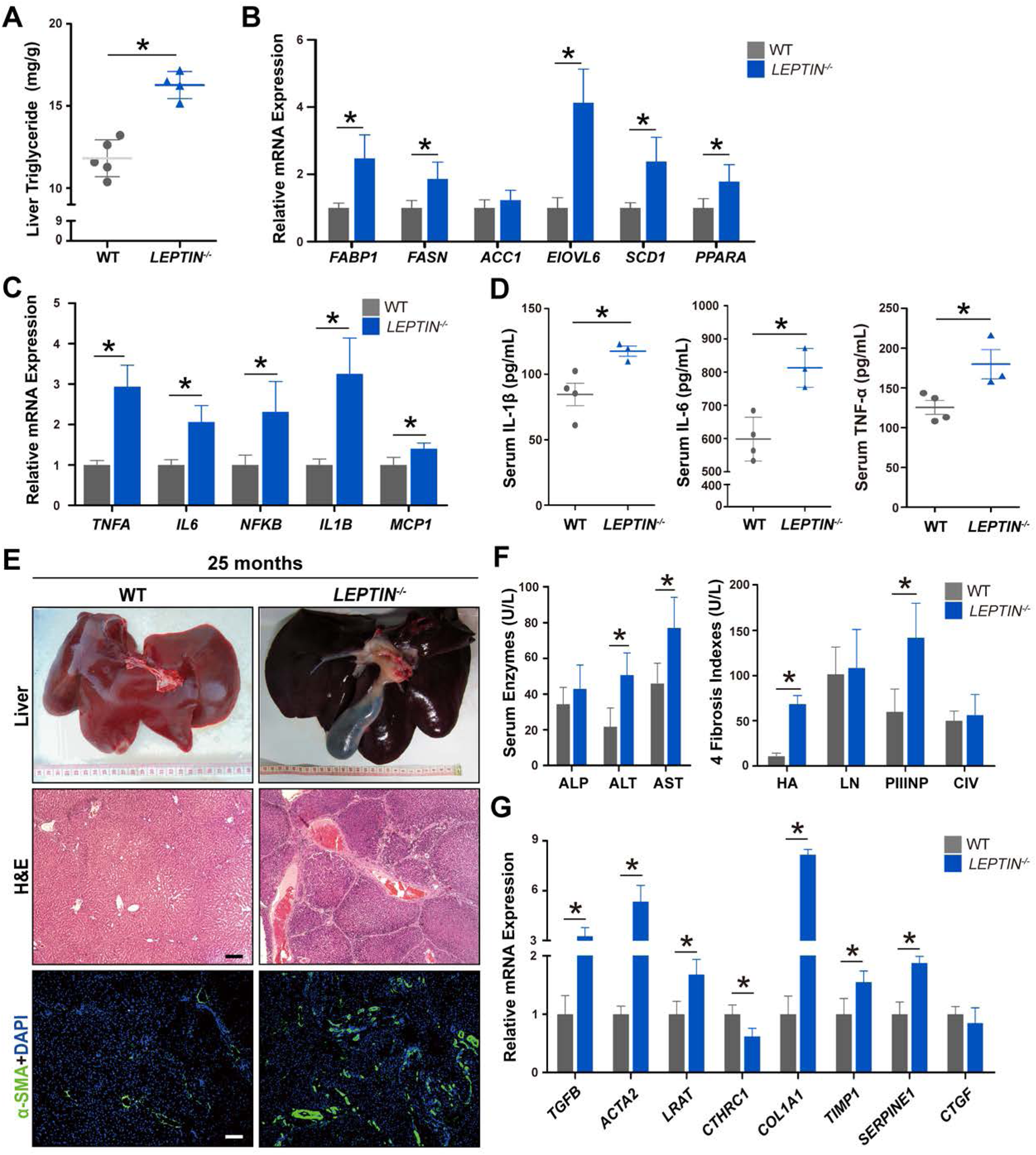
Pathological and molecular detection of liver injury in *LEPTIN^-/-^* pigs. Triglyceride concentrations **(A)** and expression of genes related to FFA synthesis **(B)** in *LEPTIN* mutant and WT pigs. Expression of inflammation related genes in liver **(C)** and the concentration of inflammatory cytokines, IL-1β, IL-6 and TNF-α in serum **(D)**. N=3-5/group. **E.** Gross images, H&E and immunofluorescent staining for α-SMA in 25-month-old *LEPTIN* mutant and WT livers (bar=20μm). **F.** Liver functional status and expression of liver fibrosis markers in serum. **G.** Expression of genes related to fibrogenesis. N=3-5/group. The bars represent the mean±SD; *P<0.05; NS, non-significant.

Liver fibrosis, classified as the third stage of non-alcoholic liver injury, is the last step of the reversible damage process observed clinically. *LEPTIN^-/-^* pig livers obtained from animals between 22 to 35 months of age demonstrated a high degree of hepatomegaly with the presence of surface fiber bundles and granule (Figure 2D). H&E staining demonstrated that the overall lobule structure was disorganized, and the fibrous septum of the portal area was significantly widened with distinct bridging lesions occurring in the portal and central vein areas in *LEPTIN^-/-^* pig livers (Figure 2D & Figure 3E). Meanwhile, Sirius Red staining revealed collagen accumulation around the hepatic lobule sinus and veins in *LEPTIN^-/-^* pig livers (damaged individuals/total individuals=3/7, 42.9%) (Figure 2D). Immunofluorescence staining demonstrated that the expression of α-SMA, a marker of fibrosis, was significantly up-regulated in *LEPTIN^-/-^* livers (Figure 3E). In the clinical diagnosis of liver fibrosis, blood indexes are used to stage the disease. In this study, the fasting blood samples were collected from *LEPTIN^-/-^* and WT pigs. The results showed that ALT and AST levels in *LEPTIN^-/-^* blood were greatly increased, indicating significant liver damage. However, comparing with WT, the level of ALP remained unchanged (Figure 3F). The levels of HA and PCIIINP in *LEPTIN^-/-^* serum were significantly elevated, indicating that fibrogenesis had occurred, although the levels of LN and CIV demonstrated no differences between *LEPTIN^-/-^* and WT livers (Figure 3F). These suggest that the severity of the fibrotic lesions in *LEPTIN^-/-^* livers was most likely in the middle or advanced stage of fibrosis but not cirrhosis. The expression of fibrotic markers, *TGFΒ*, *ACTA2*, *LRAT* and *COL1A1*, were increased greatly in *LEPTIN^-/-^* livers (Figure 3G). Previous studies had shown that *CTHRC1* limits collagen deposition (13); *TIMP1* inhibits the degradation of the main components of the extracellular matrix (14); and *SERPINE1* inhibits the dissolution of fibrinolysis (15). The reduction of *CTHRC1* and elevation of *TIMP1* and *SERPINE1* expression helps explain the observed collagen accumulation in *LEPTIN^-/-^* livers.

Following additional H&E staining, PAS staining, Sirius red staining, Brunt’s scoring criteria and Scheuer scoring criteria (16), the stage of non-alcoholic liver inflammation and fibrosis in *LEPTIN^-/-^* pigs was determined. The statistical results showed that the average NASH score was 4.47±0.86 at 12-22 months in *LEPTIN^-/-^* pig. By 22-35 months, the average fibrosis score in *LEPTIN^-/-^* pig was 3.00±0.27. In humans, the highest score observed clinically is 4.0. Thus, the stage of fibrosis in *LEPTIN^-/-^* pigs was clinically assessed in the range of moderate and severe (Table 1).

**Table 1.**
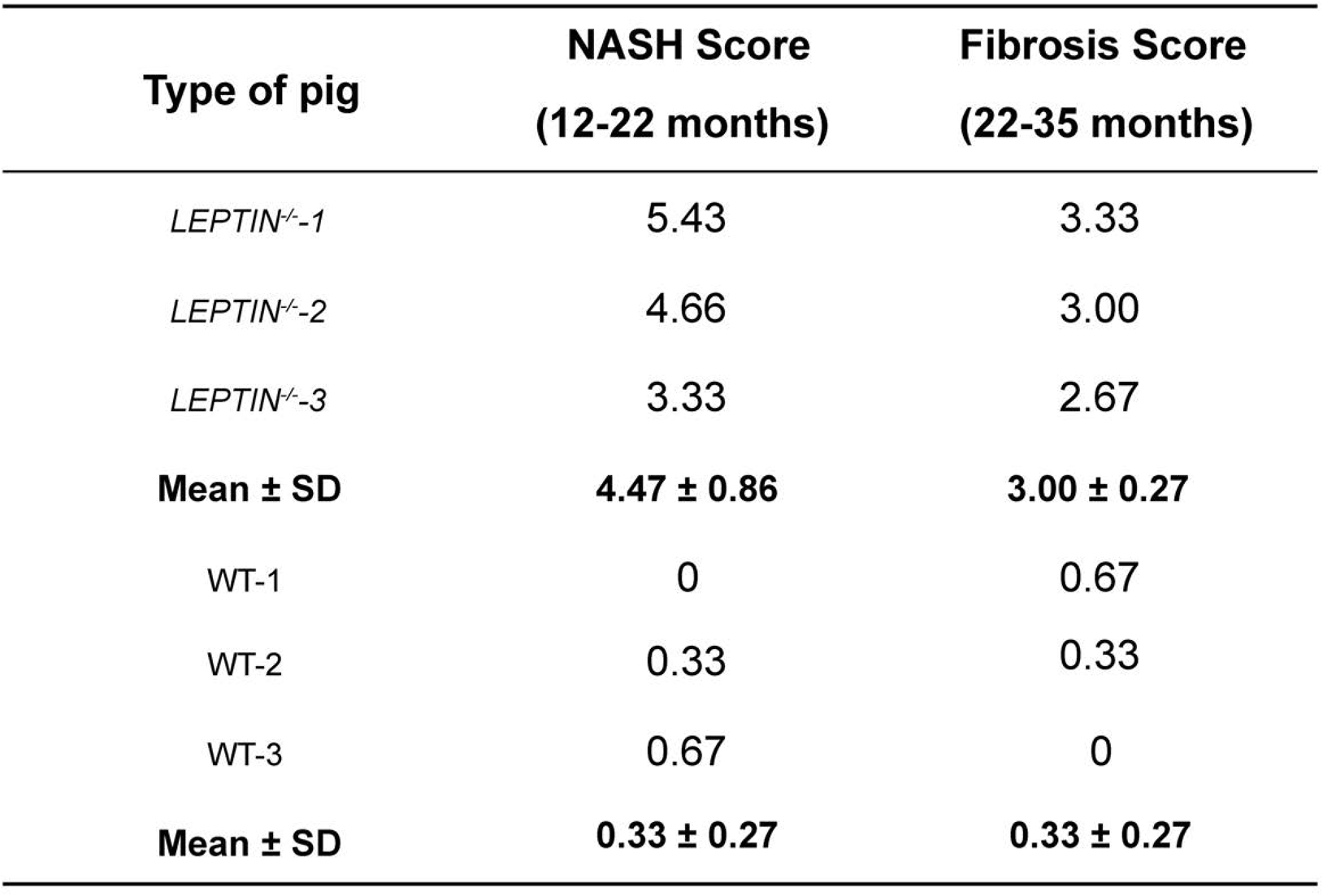
Assessment of liver fibrosis and NASH staging in pigs. Brunt’s and Scheuer scoring criteria were used to determine the stage of non-alcoholic liver inflammation and fibrosis in *LEPTIN^-/-^* and WT pigs.

### Porcine LEPTIN deficiency results in lipid peroxidation via the JAK2-STAT3 pathway

To identify the mechanism driving liver fibrosis in *LEPTIN^-/-^* pigs, several signaling pathways proven to be affected by LEPTIN were analyzed. Utilizing WB and real-time PCR analyses, no significant changes in the expression of mTOR, MAPK and PI3K-AKT pathway related proteins and genes were observed (Figure 4- figure supplement 1). Interestingly, phosphorylation of JAK2 and STAT3 was significantly reduced in *LEPTIN^-/-^* livers. In addition, altered expression of genes and proteins related to the JAK-STAT signaling pathway was also observed in response to LEPTIN deletion (Figure 4A & Figure 4- figure supplement 1B). The WB assay demonstrated that the phosphorylation of JAK2 and STAT3 was significantly reduced in *LEPTIN^-/-^* cells (Figure 4A). In particular, SOCS3, a negative regulator of cytokines and hormone transduction (17), and SREBP-1c, related to fat synthesis (18) were analyzed. STAT directly inhibits SREBP-1c, and SOCS3 promotes SREBP-1c expression (19). Both SOCS3 and SREBP-1c expression was elevated in *LEPTIN^-/-^* livers (Figure 4B). Up-regulation of ACSL3 and ACSL5 in *LEPTIN^-/-^* indicates increased synthesis of acyl-CoA. The expression of CPT1a and CPT2 were also up-regulated in *LEPTIN^-/-^* livers, suggesting that FFA oxidation was activated. Immunofluorescence staining of SOCS3, SREBP1c and ACSL3 demonstrated increased expression in *LEPTIN^-/-^* fibrotic livers, suggesting that FFA synthesis was enhanced. In addition, the increased CPT1a and CPT2 expression suggested that acyl-coA transportation from the endoplasmic reticulum to mitochondria was enhanced (Figure 4- figure supplement 2).

**Figure 4.**
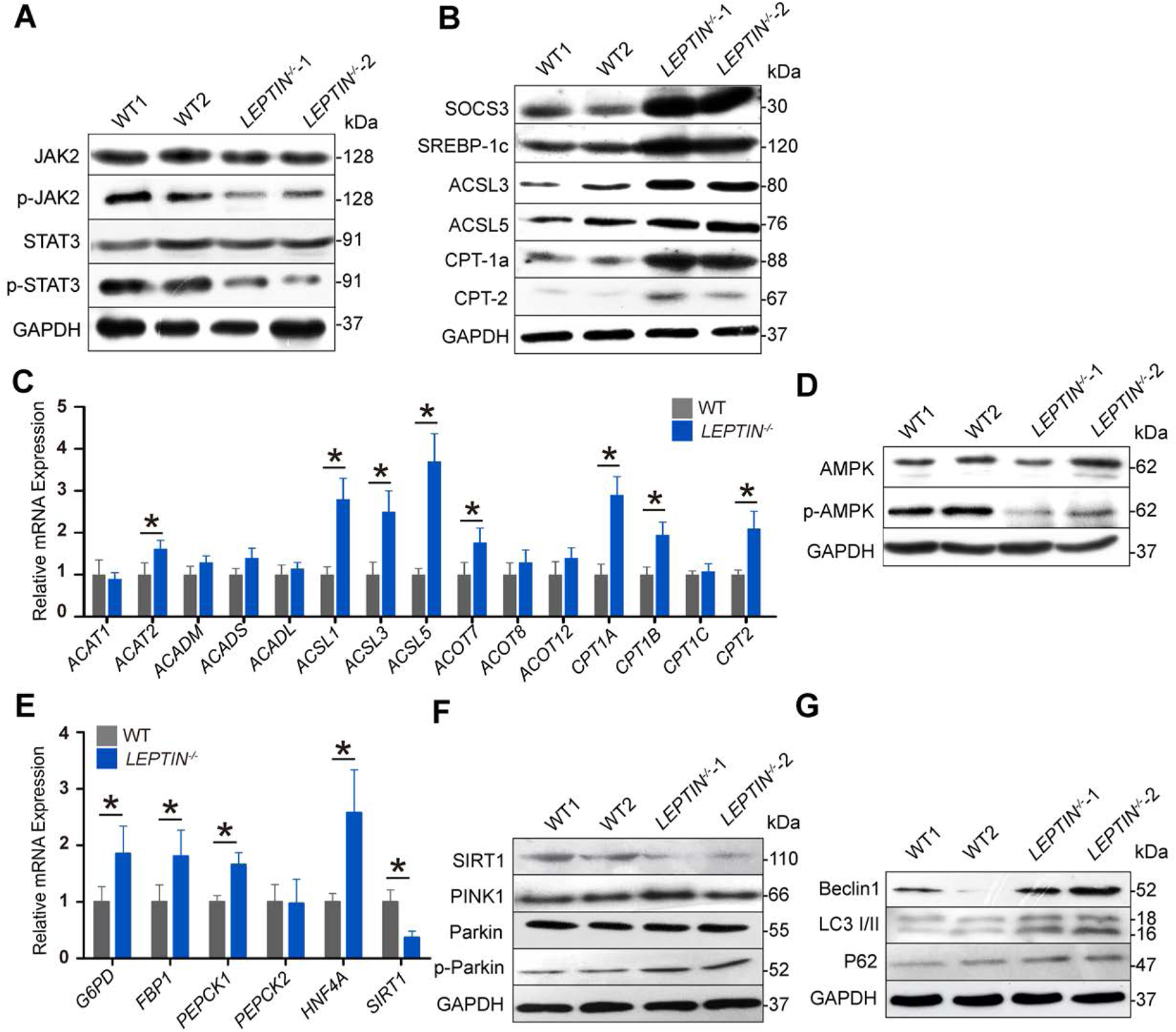
Alteration of JAK-STAT and AMPK pathway related genes in *LEPTIN^-/-^* pig livers. **A.** WB analysis of JAK-STAT signaling proteins. **B.** WB analysis of FFA synthesis and oxidation related proteins. **C.** Expression of β-oxidation related genes. **D.** WB analysis of AMPK signaling proteins. **E.** Expression of gluconeogenesis related genes. **F.** WB analysis of SIRT1-mediated autophagy related proteins. **G.** WB analysis of mitochondrial autophagy markers. N=3/group. The bars represent the mean±SD; *P<0.05.

Since LEPTIN deficiency affects β-oxidation, genes involved in the oxidation process were analyzed: *ACAT1* and *ACAT2* are acyl-coA transferases; *ACADM, ACADS* and *ACADL* are acyl-coA dehydrogenases; *ACSL1, ACSL3* and *ACSL5* are acyl-coA synthetizes; *ACOT7, ACOT8* and *ACOT12* are acyl-coA thioesterases; *Cpt1a, Cpt1b, Cpt1c* and *Cpt2* are carnitine transferases. Among them, *ACAT2*, *ACSL1*, *ACSL3*, *ACSL5*, *ACOT7*, *CPT1A*, *CPT1B* and *CPT2* were significantly up-regulated. This suggested that β-oxidation has mainly been strengthened during acyl-coA synthesis and transfer to the mitochondria (Figure 4C).

### LEPTIN deficiency enhances gluconeogenesis and mitochondrial autophagy through the AMPK pathway in pigs

Gluconeogenesis disorder is the most typical feature of insulin resistance in which HNF-4α synergizes with PGC-1α, FOXO1 and other key enzymes to induce gluconeogenesis (20), and Leptin regulates the AMPK pathway by inhibiting HNF-4α and promoting SIRT1 expression (21). In addition, PEPCK and G6Pase are speed limiting enzymes of glycogenesis (22). Based on the conclusions of these previous studies, protein or gene expression levels of these markers in biological pathways or biochemical processes in *LEPTIN^-/-^* pig livers were detected. It confirmed that the lack of Leptin down-regulated phosphorylation of AMPK due to increased Hnf-4α and decreased SIRT1 expression in *LEPTIN^-/-^* pig livers. PEPCK and G6Pase were also significantly up-regulated (Figure 4D&E). In addition, the fluorescence staining results demonstrate increased PEPCK and G6Pase expression with highly expressed α-SMA in *LEPTIN^-/-^* fibrotic livers, especially in the portal and perivenous areas (Figure 4- figure supplement 3).

SIRT1 regulates mitochondrial autophagy and proliferation through PINK1/Parkin and PGC-1α/TFAM, respectively (23). Initially markers of mitochondrial autophagy, Beclin1, LC3-II and P62 (24) were examined. The expression levels of PINK1 and phosphorylation of PARKIN were largely increased (Figure 4F), and the expression of Beclin1, LC3-II and P62 were also significantly increased in *LEPTIN^-/-^* livers (Figure 4G). In contrast with the autophagy related genes, most of the mitochondrial synthesis related genes were not altered in *LEPTIN^-/-^*. No expression changes for the mtDNA genes *ATP6, COX1* and *ND1* were observed via qPCR (Figure 4- figure supplement 4A). mtDNA copy number was calculated using Gcg as the internal reference, and the mtDNA copy number was not altered in *LEPTIN^-/-^* pig livers (Figure 4- figure supplement 4B), suggesting that the down-regulation of SIRT1 in *LEPTIN^-/-^* pig livers enhances mitochondrial autophagy but does not affect mitochondrial synthesis.

### *LEPTIN^-/-^* pigs undergo oxidative stress

Both lipid peroxidation and mitochondrial autophagy produce excessive ROS which in turn causes oxidative stress (25). Heatmap analysis based on DEGs and expression difference fold change was performed using the Omicshare software (Figure 5- figure supplement 1). GO enrichment analysis identified oxidation-reduction GO terms enriched for DEGs (Figure 5- figure supplement 2). In addition, the CYP2 mediated arachidonic acid metabolism and exogenous toxic substances metabolic pathways were unexpectedly enriched for DEGs in *LEPTIN^-/-^* pig livers (Figure 5- figure supplement 3 & Table. S2). Since the CYP2 enzyme mediates oxidative stress (26), real-time PCR was utilized to monitor the expression of CYP2 family genes (Figure 5A). The level of CYP2E1 in *LEPTIN^-/-^* liver was nearly 3 times higher than in WT (Figure 5B). The SOD enzyme activity was significantly reduced in *LEPTIN^-/-^* livers and sera, which could contribute to the excessive accumulation of ROS. Meanwhile, NO was up-regulated, and its reaction with OH increased the production of toxic ROS, which in turn led to increased MDA (Figure 5C-E). These data indicate that LEPTIN deletion enhances oxidative stress in pig livers.

**Figure 5.**
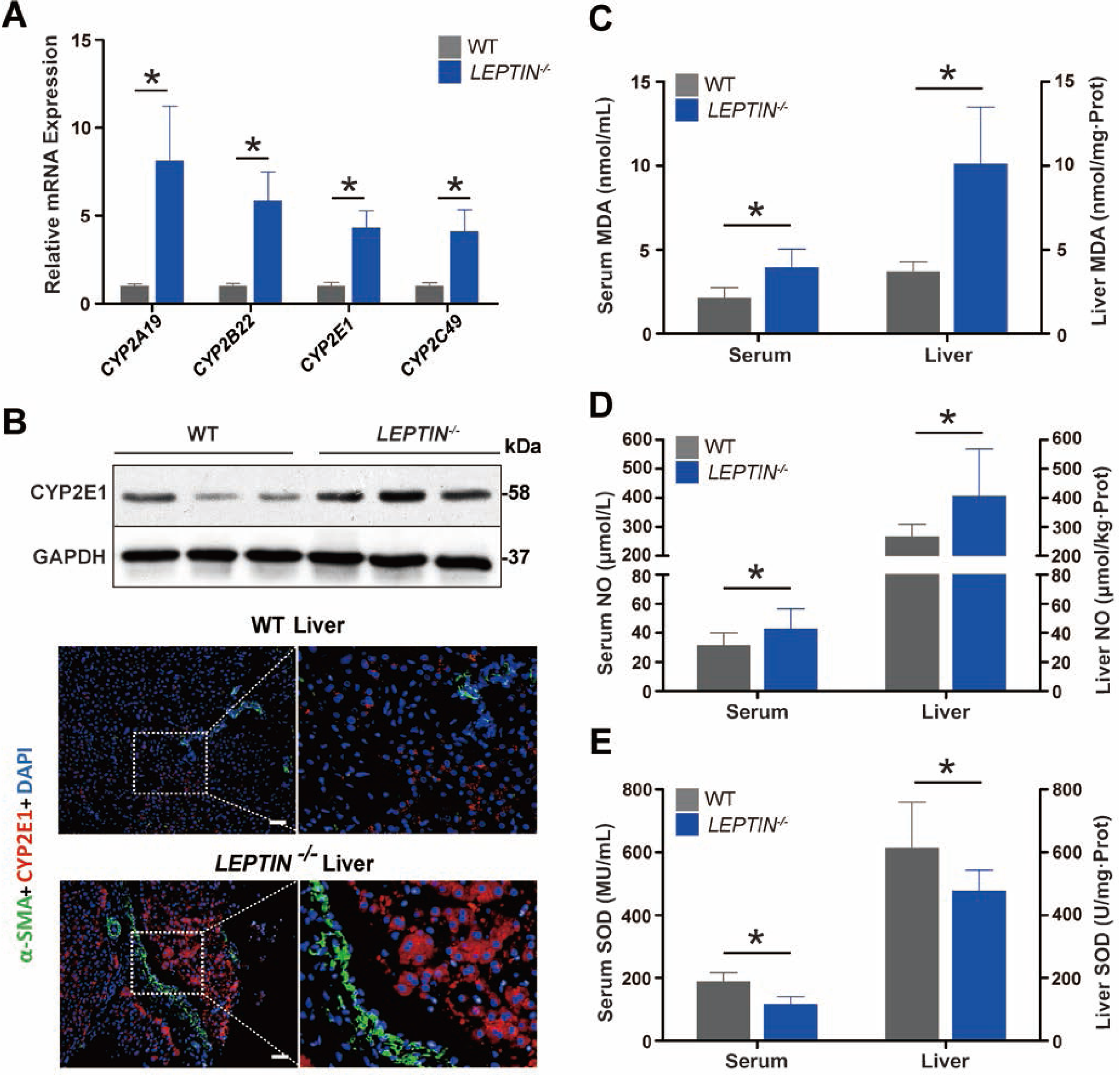
Detection of oxidative stress in *LEPTIN^-/-^* pig livers. **A.** Expression analysis of CYP2 family genes by real-time PCR. **B.** WB analysis and Immunofluorescence staining of CYP2E1. Scale bar represents 100μm. **C-E.** Detection of oxidative stress markers in the liver and serum. Content of MDA **(C),** NO **(D)** and SOD **(E)**. N=3/group. The bars represent the mean±SD; *P<0.05.

### LEPTIN deficient rat livers are void of hepatic fibrosis, mitochondrial autophagy and oxidative stress

Interestingly, published studies and data presented in this study demonstrate that, in contrast with pigs, Leptin deficiency in rodents does not result in hepatic fibrosis (27, 28). Compared with WT rats, the *Leptin^-/-^* rats have larger body and liver sizes (Figure 6A). H&E staining of *Leptin^-/-^* rat livers revealed fatty liver, lipid degeneration and balloon-like degeneration (Figure 6B). Sirius red staining, α-SMA staining and the four serum hepatic fiber indicators analysis showed no obvious collagen accumulation and fibrogensis occurring in *Leptin^-/-^* rat livers (Figure 6B-C). Furthermore, testing for SOD, MDA and NO found no obvious differences between *Leptin^-/-^* and WT rat livers (Figure 6- figure supplement 1A-C). This suggests that oxidative stress does not occur in *Leptin^-/-^* rat livers. WB and immunofluorescence staining revealed that the level of CYP2E1 was lowered (Figure 6D & Figure 6- figure supplement 1D). To investigate whether Leptin deletion in rats alters the JAK2-STAT3 pathway, p-JAK2/JAK2, p-STAT3/STAT3, SREBP-1C, SOCS3, CPT-1A and CPT-2 expressions were analyzed by WB and immuno-fluorescence staining. As observed in the *LEPTIN^-/-^* pig liver, the level of p-JAK2 and p-STAT3 were decreased compared to WT rat livers (Figure 6- figure supplement 2A) whereas the SREBP-1C and SOCS3 were significantly increased in *Leptin^-/-^* rat livers (Figure 6- figure supplement 2B&C), suggesting that the FFA synthesis was activated in *Leptin^-/-^* rat liver. However the expression of CPT-1A and CPT-2 in *Leptin^-/-^* rat livers were not altered (Figure 6- figure supplement 2B&C).

**Figure 6.**
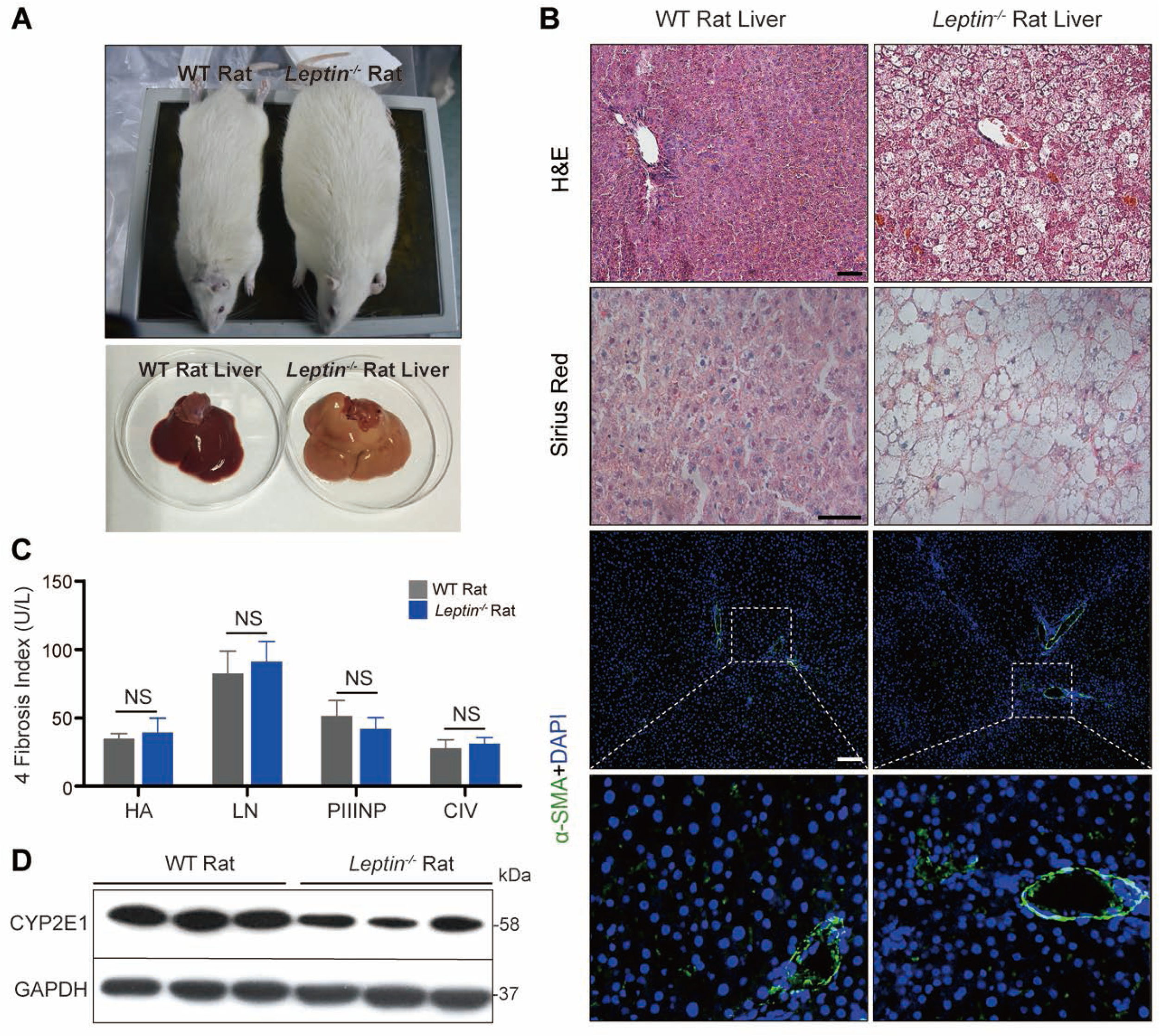
Analysis of liver injury and related signaling pathways in *Leptin^-/-^* rats. **A.** Comparison of *Leptin^-/-^* and WT rat bodies and livers. **B.** H&E, Sirius red and α-SMA staining analysis of rat livers. Scale bar represents 100μm. **C.** Detection of fibrosis markers in serum.

Previous studies in pigs have demonstrated that Leptin deficiency promotes mitochondrial autophagy, representing another major source of oxidative stress. Therefore, mitochondrial autophagy related markers were examined. Beclin1, LC3-Ⅱ and P62 expression showed no significant differences between *Leptin^-/-^* and control rats (Figure 6- figure supplement 3A). It was interesting to note that, in contrast with *LEPTIN^-/-^* pigs, SIRT1 expression in *Leptin^-/-^* rat liver was up-regulated (Figure 6- figure supplement 3B). This is quite different from the results in pigs and may be a key reason why *Leptin^-/-^* rat livers do not have mitochondrial autophagy.

In order to investigate the underlying causes of this phenomenon of opposite trends in protein expression between the two animals, WB was performed to detect the phosphorylation of AMPK pathway related genes. Interestingly, in contrast with *LEPTIN^-/-^* pig, the phosphorylation of AMPK was significantly increased in *Leptin^-/-^* rats (Figure 6- figure supplement 3C), providing a mechanism through which SIRT1 expression declined as SOCS3 and SREBP-1C expression increased.

## Discussion

The liver closely communicates with adipose tissue and liver injury is observed in nearly 80% of obese patients (29). Since mechanistic studies of liver fibrogenesis are difficult to conduct in patients, current knowledge of obesity-induced liver fibrosis is mainly derived from Leptin-deficient animal models. The choice of Leptin is based on its important roles in mediating energy homeostasis. Serum Leptin concentrations demonstrate an association with NAFLD which is mediated through insulin secretory dysfunction and insulin resistance in obese patients (30). NAFLD, the hepatic manifestation of the metabolic syndrome, comprises steatosis, steatohepatitis, hepatic fibrosis, cirrhosis and hepatocellular carcinoma (31). However, both previously published studies and work presented here suggests that two of the most commonly used rodent models, *ob/ob* mouse and Zucker rat, do not develop fibrosis observed in NAFLD patients. In an effort to develop a model that displays fibrogenesis as observed by patients, a *LEPTIN^-/-^* pig model was created. Consistent with human clinical data, the *LEPTIN^-/-^* pigs demonstrated obstructed body glucose regulation leading to insulin resistance and type II diabetes by 12-months of age. Interestingly, around the same age (6-12 month), 60% of *LEPTIN^-/-^* pig livers displayed lipid deposition and hepatocyte steatosis. The severity of liver injury also worsened with age in *LEPTIN^-/-^* pigs. Lobular inflammation and hepatocyte ballooning appeared in 33% *LEPTIN^-/-^* of pig livers between 12-22 months, whereas 42.9% *LEPTIN^-/-^* pig livers developed various degrees of fibrosis by 22-35 months of age (Figure 2). The development of liver injury in *LEPTIN^-/-^* pigs mirrored the pathological progression of NAFLD observed in obese patients. *LEPTIN^-/-^* pigs appeared to closely model the process of fibrogenesis observed in patients, which supports their use for investigation of the mechanisms underlying fibrogenesis in patients.

The mechanisms leading to NAFLD are unclear to date. In obese subjects, FFAs seem to be misrouted to ectopic sites like hepatic tissues, resulting in steatosis. Steatosis induces the production of pro-inflammatory mediators like TNF-α, IL-6 and IL-1β. These cytokines promote the recruitment and activation of Kupffer cells, which induce inflammation and hepatic insulin resistance via SOCS3. The accumulation of fat in the liver leads to lipotoxicity and dysfunctional mitochondria, which further causes oxidative stress due to imbalanced ROS production and protective oxidants, eventually leading to hepatocyte death in NAFLD patients. The reduced phosphorylation of JAK2 and STAT3 in LEPTIN deficient pig livers led to hepatic insulin resistance and increased fat synthesis marked by activation of SOCS3 and SREBP-1c, respectively, further confirms the value of the pig model (Figure 4). This data is consistent with observed alterations in the JAK/STAT pathways in the pathogenesis of metabolic disease (32). The excess intracellular FFAs observed in *LEPTIN^-/-^* pigs resulted in intrinsic endoplasmic reticulum stress. The toxic pathways were represented through the generation of ROS and increased mitochondrial β-oxidation, which further caused hepatocyte death and fibrosis. Interestingly, in contrast with pigs, although the Leptin deletion in rats altered the JAK2 and STAT3 phosphorylation and caused fatty liver, the level of damage was not severe enough to drive mitochondrial β-oxidation and oxidative stress, which may explain why fibrosis did not occur in *Leptin^-/-^* rat livers (Figure 6).

The progression of NAFLD to NASH has been characterized as a mitochondrial disease occurring at an early human disease stage (33). Lipotoxicity encompasses the dysregulation of the intracellular lipid composition, resulting in mitochondria dysfunction, stimulating ROS production, oxidative stress, and impaired FFA oxidation (34, 35). AMPK is a heterotrimeric enzyme which regulates cell growth, proliferation, autophagy and apoptosis (36). In NAFLD, activation of AMPK in the liver inhibits the synthesis and oxidation of FFAs, leading to the reduction of ectopic lipid accumulation and improved insulin action (37). The current study on *Leptin^-/-^* rats provided evidence that AMPK activation might be crucial for prevention of liver damage. This is further supported by the observed inactivation of the AMPK pathway in *LEPTIN^-/-^* pigs, which drove mitochondrial autophagy, hepatic cell death and fibrosis. Whether LEPTIN is required for activation of the AMPK pathway, or AMPK mediated autophagy appears as a defensive reaction to liver injury has not been previously addressed in the pig. Nevertheless, one collective conclusion from our study is that the activity of AMPK represents a potential predictive biomarker of the severity of liver injury. Activation of AMPK using pharmacological agents may also represent a potential therapeutic avenue for preventing the progress of NASH to fibrosis.

Liver fibrosis is defined by the excessive accumulation of extracellular matrix (ECM) proteins. Activated hepatic stellate cells (HSCs) in the liver are the major source of collagen production, leading to an imbalance in the formation and degradation of the ECM. Activation of HSCs in the injured liver is regulated by fibrogenic and pro-inflammatory cytokines (38). Recent evidence demonstrated that adiponectin induces apoptosis and inhibits activation of HSCs through the AMPK pathway (39). Thus, the reduced AMPK signaling observed in *LEPTIN^-/-^* pig livers may promote the activation of HSCs, which in turn induce the deposition of ECM proteins and development of fibrosis; whereas the activated AMPK in *Leptin^-/-^* rat livers may lead to the apoptosis of HSCs, preventing the accumulation of ECM proteins.

The underlying mechanisms by which NASH transitions to fibrosis are not completely understood. Hepatic fibrosis occurs in 40-50% of patients with NASH and approximately 30-40% of NAFLD patients develop NASH. It has been estimated that the hepatic fibrosis stage is the strongest predictor of mortality in NAFLD patients. The present understanding of the dynamics in NAFLD progression has emerged from genetically modified rodents, such as *ob/ob* mice and Zucker rats. These animals are universally used in obesity and diabetic research. However, they fail to develop hepatic fibrosis. The LEPTIN deficient pigs generated in this study mirror the progression of hepatic fibrosis observed in NAFLD patients. Loss of LEPTIN in pigs led to β-oxidation and oxidative stress and, in combination with AMPK mediated mitochondrial autophagy, increased liver fibrosis (Figure 7). In sum, LEPTIN-deficient pigs provide an ideal model to investigate the full spectrum of human NAFLD and develop new strategies for the diagnosis and treatment of NAFLD.

**Figure 7.**
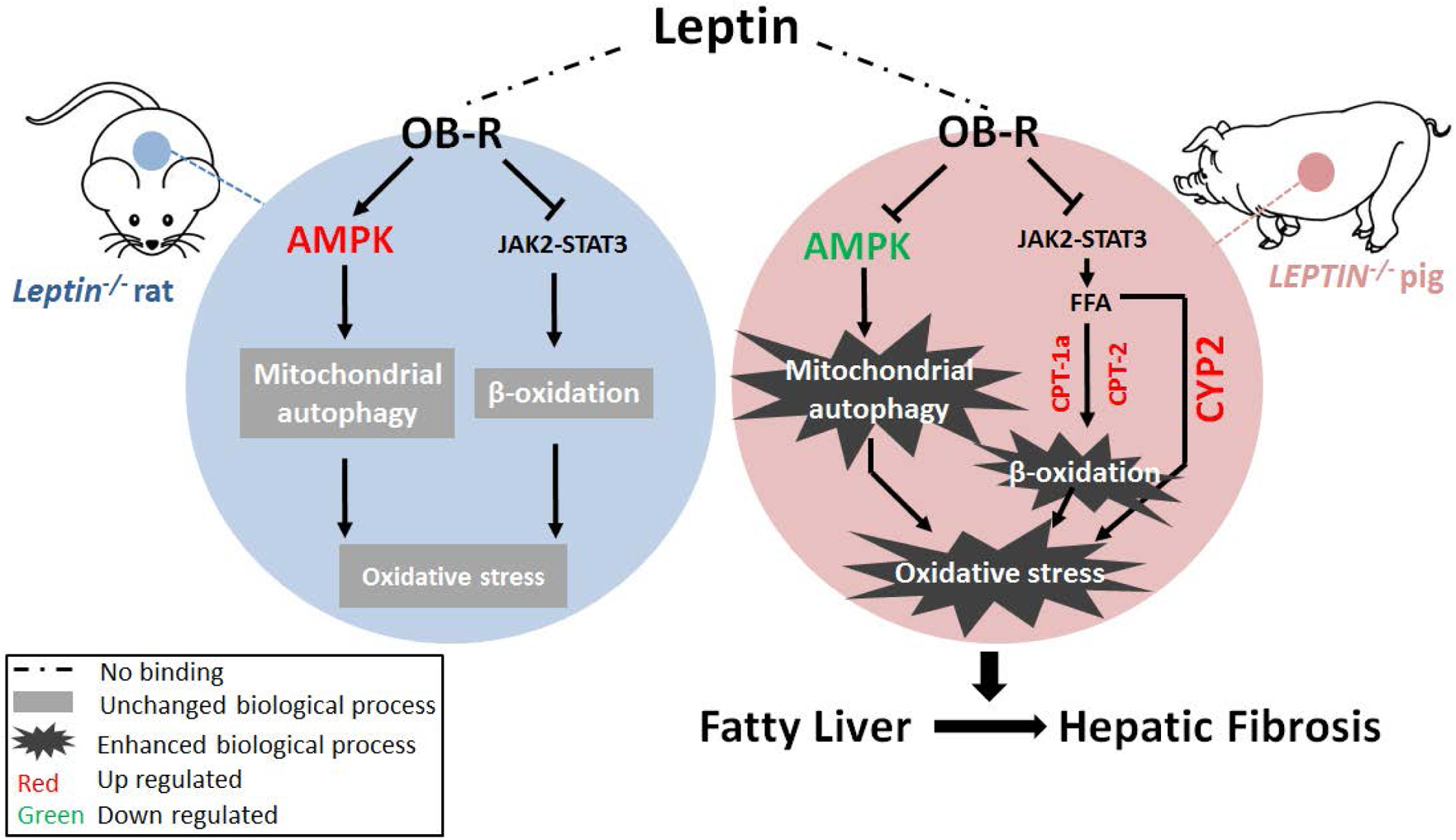
Proposed mechanistic model of fibrosis progression in *LEPTIN^-/-^* pig liver. Reduction of p-AMPK promotes mitochondrial autophagy. Meanwhile, activation of JAK2/STAT3 signaling pathway leads to enhanced FFA oxidation and oxidative stress. Oxidative stress drives the development liver fibrosis in *LEPTIN^-/-^* pigs.

## Materials and Methods

Animal experiments were approved by the Animal Care Committee of the China Agricultural University (Plan book number SKLAB-2016-84) and performed according to the Chinese Animal Welfare Act. All chemicals used were purchased from Sigma-Aldrich Co. (Alcobendas, Madrid, Spain) unless otherwise indicated.

### Experimental animals

The pig species used to generate *LEPTIN^-/-^* pigs was the Chinese experimental mini-pig (10). Both wild type and *LEPTIN^-/-^* pigs were provided with same manufactured diet. The cereal grain in dry form has been ground to supply to the pigs using an ad libitum feeding at fixed time (7:00-8:00, 1.75kg max; 12:00-13:00, 1.25kg max; and 17:00-18:00, 1kg max) every day. As shown in Supplemental Table 5, the nutrition content of pig diets were changed with age. The *Leptin^-/-^ r*ats were generated by the Sen Wu laboratory (11). Both wild type and *Leptin^-/-^* rats were provided with the same always abundant manufactured diet (Table S5).

### Generation of *LEPTIN^-/-^* pigs

Zinc finger nuclease (ZFN) plasmids targeting the *LEPTIN* gene were designed by Sigma-Aldrich. Fetal fibroblasts were isolated from 30-day old male pig embryo. Approximately 1×10^6^ CEMP fetal fibroblasts were transfected with 2μg of each ZFN vector. Following G418 selection (500μg/mL, Promega), positive clones were collected for somatic cell nuclear transfer. PCR was performed following by Sanger sequencing to confirm the genetic sequence of the *LEPTIN* target region (primer sequence : Forward 5’-GATTGTGTGGAAGGGGAAGA-3’, Reverse 5’-GAGGTCCGCACAGGCTCTC T-3’.

### Western blotting (WB)

Total protein of adipose tissue, liver tissue, hypothalamus tissue and serum were obtained from the mutant and wild-type pigs. Electrophoretically separated proteins were detected by specific primary antibodies (Table S3).

### Enzyme linked immunosorbent assay (ELISA)

Serum was isolated from blood collected from pigs after fasting overnight. ELISA kit was used to detect porcine IL-1β, IL-6 and TNF-α (Dan Shi Biological Technology, Shanghai).

### Growth parameters and body composition

Body weight of *LEPTIN^-/-^* and control pigs were examined weekly or monthly. The percentage of total body fat was determined by using multi-slice computed tomography (MSCT) with 64 rows, 128-slice MSCT scanner (Siemens). MSCT was performed with the following parameters: 120kV, 500ms, and 1.5mm splice thickness. The 3D reconstruction and analysis of data were performed by Sygo Fastview software.

### Assessment of clinical indicators of liver disease

Triglycerides, insulin, cholesterol, HDL, LDL, AST, ALT, ALP and four indexes of hepatic fibrosis were analyzed by the Beijing CIC Clinical Laboratory, China Agricultural University Veterinary Teaching Hospital and the endocrinology department of Beijing Tongren Hospital.

### HOMA model assessing type II diabetes

Homeostatic model assessment (HOMA) included the following values, calculated on the basis of two parameters of fasting plasma glucose and insulin: HOMA-IR=(FPG, mmol/L)×(FINS, mIU/L)/22.5, assessment of insulin resistance; HOMA-IS=1/(FPG, mmol/L)×(FINS, mIU/L), assessment of insulin sensitivity; HOMA-β=20×(FPG, mmol/L)/[(FPG, mmol/L)-3.5] (%), assessment of islet β cell function. FPG, Fasting plasma glucose and FINS, fasting insulin.

### Intravenous glucose tolerance test (IVGTT)

After 12-15 hours of fasting, pig blood samples were collected from the anterior chamber vein. 50% glucose solution was injected into the ear vein with a dose of 1.2mL/kg. Glucose concentrations were measured at 0 min (before injection), 1 min, 5 min, 10 min, 15 min, 30 min, 60 min and 90 min (after injection).

### Histological analysis

Adipose, liver and pancreatic tissues were fixed in 10% neutral buffered formalin solution and paraffin-embedded. 5μm-thick sections were used for hematoxylin-eosin (H&E) analysis, oil red O dye detection, periodic acid schiff (PAS) analysis (Beijing Solarbio life science), and Sirius red (Beijing Solarbio life science) analysis. α-SMA (ab5694, abcam) antibody was used for immunofluorescence using the ABC method (Vector Laboratories).

### RNA isolation and quantitative RT-PCR

Total RNA was isolated from pig fat or liver tissues by TRIzol and RNeasy Mini Kit (QIAGEN). Then RNA samples were reverse transcribed to cDNA by M-MLV reverse transcription kit (Promega). The levels of relevant mRNAs were quantitated by real-time PCR using One Step SYBR GREEN RT-PCR Kit (Roche) in a Light Cycler instrument (Roche Applied Science, Mannheim, Germany). The specific primers for related genes were designed by Primer3 (v.0.4.0), PrimerBank, primer premier5.0 and NCBI BLAST (Table S4).

### Detection of oxidative stress markers

Blood samples were collected from pigs after fasting overnight. Liver tissue was homogenized in cold 0.9% normal saline. The determination of NO requires the use of a specific homogenate. The total SOD activity detection kit (WST-8 method) (Beyotime, S-0101), lipid oxidation (MDA) assay kit (Beyotime, S-0131) and nitric oxide detection kit (Beyotime, S-0021) were utilized following the manufacturer’s instructions.

### Transcriptome analysis

The total liver RNA of three *LEPTIN^-/-^* and WT pigs from 22-35 month of age was extracted. The library construction and sequencing were performed with the steps of purifying mRNA, interrupting mRNA, synthesizing cDNA, selecting fragments, and PCR amplification. The qualified libraries were generated on Illumina Cbot for cluster generation, and then Illumina HiSeqTM2500 was used for transcriptome Sequencing. The purity, concentration and integrity of RNA samples were measured by Nano Drop, Qubit 2.0 and Agilent 2100 before sequencing. The cDNA library was constructed using the Illumina TruseqTM RNA kit(Illumina, USA). The sequencing read length is PE125. Nearly 50GB per sample of raw data was obtained by sequencing, and 6-8GB clean data of each sample was obtained after removal of reads containing low sequencing quality with connectors and duplicates. After clean reads were obtained, HISAT was used for sequence alignment with the reference genome (Sus_scrofa. Sscrofa10.2.dna.chromosome) to obtain the information of the reference genes and the Mapped reads were obtained. By using the Cuffdiff component of Cufflinks software, the gene expression levels were quantified. Pearson’s correlation co-efficient (r^2^) was used as an indicator to evaluate the biological correlation of repetition. Then FPKM was used to determine the expression abundance of transcripts. The absolute value of Log2 Fold Change ≥1, and the corrected P value (FDR) < 0.05, was taken as the key index to screen the differential genes. GO and KEGG software were used for differential genes enrichment and signaling pathways screening. In GO and KEGG analysis, P < 0.05 was used as the selection criteria. DAVID and KOBAS software were used for gene function analysis. The RNA-Seq data was deposited in Gene Expression Omnibus (GEO) under the accession number GSE176023.

### Statistical analysis

The experimental data are presented as the mean ± SD and were analysed by paired Student’s *t* test using SPSS15.0 software to compare the mutant and wild-type pigs. *P* value <0.05 was considered significant.

### Data availability

The datasets during and/or analysed during the current study available from the corresponding author on reasonable request.

## Acknowledgments

We are grateful for the gift of *Leptin^-/-^* rats by Dr. Sen Wu. We thank Dr. Zuoxiang Liang for the technical support. This work is supported by National Natural Science Foundation of China 31972566, National Genetically Modified Organisms Breeding Major Projects of China 2016ZX08009003-006, National Basic Research Program 2016YFA0100202.

## Author contributions

T.T., Z.S., R.W., S.J., and Q.W. performed the experiments; X.H., and N.L. analyzed the data; Y.X. conceived and designed the experiments, and wrote the manuscript with input from all authors. All authors have read and approved the manuscript.

## List of abbreviations

ACADL: acyl-CoA dehydrogenase long chain
ACADM: acyl-CoA dehydrogenase medium chain
ACADS: acyl-CoA dehydrogenase short chain
ACAT1/2: acetyl-CoA acetyltransferase 1/2
ACOT7/8/12: acyl-CoA thioesterase 7/8/12
ACSL1/3/5: Acyl-CoA synthetases 1/3/5
ACTA2: Alpha-actin 2
ALP: alkaline phosphatase
ALT: alanine aminotransferase
AMPK: AMP-activated protein kinase
AST: aspartate aminotransferase
ATP6: mitochondrial ATPase subunit 6
CEMP: Chinese miniature experiment pig
CIC: China isotope corporation
CIV: collagen type IV
COL1A1: collagen Type I Alpha 1
COX1: mitochondrial Cytochrome c oxidase subunit I
CPT 1A/1B/2: carnitine palmitoyltransferase 1A/1B/2
CTHRC1: collagen triple helix repeat containing 1
DEGs: differentially expressed genes
ECM: extracellular matrix
ELISA: enzyme linked immunosorbent assay
ELOVL6: elongaseofverylongchainfattyacids 6
FABP1: fatty acid binding protein
FASN: fatty acid synthetase
FDR: false discovery rate
FFA: free fatty acid
FINS: fasting plasma insulin
FPG: fasting plasma glucose
G6Pase: glucose-6 phosphatase
GAPDH: glyceraldehyde-3-phosphate dehydrogenase
GO: gene ontology
H&E: hematoxylin-eosin
HA: hyaluronic acid
HDL: high density lipoprotein
HISAT: hierarchical indexing for spliced alignment of transcripts
HNF-4α: hepatocyte nuclear factor 4-alpha
HOMAs: Homeostasis model assessment
HSCs: hepatic stellate cells
IL-1β/6: interleukin 1 beta/6
IR: insulin resistance
IS: insulin sensitivity
IVGTT: intravenous glucose tolerance test
JAK2: janus kinase 2
KEGG: kyoto encyclopedia of genes and genomes
LC3: microtubule associated protein 1 light chain 3
LDL: low density lipoprotein
LN: laminin
LRAT: lecithin retinol acyltransferase
MCP-1: monocyte chemotactic protein 1
MDA: malondialdehyde
MSCT: multi-slice computed tomography
NAFLD: non-alcoholic fatty liver disease
NASH: non-alcoholic steatohepatitis
ND1: mitochondrial NADH-ubiquinone oxidoreductase chain 1
NFKB: nuclear factor kappa-light-chain-enhancer of activated Bcells
NO: nitric oxide
PAS: periodic acid schiff
PEPCK: phosphoenolpyruvate carboxylase
PGC-1α: peroxisome proliferator activated receptor gamma coactivator 1 alpha
PIIINP: procollagen-III-peptide
PINK1: phosphatase and tensin homolog induced kinase 1
PPARA: peroxisome proliferator-activated receptor alpha
ROS: reactive oxygen species
SCD1: stearoyl-CoA desaturase 1
SERPINE1: serpin family E member 1
SIRT1: sirtuin 1
SOCS-3: suppressor of cytokine signaling-3
SOD: superoxide dismutase
SREBP-1c: sterol-regulatory element-binding protein-1c
STAT3: signal transducer and activator of transcription 3
TFAM: transcription factor A, mitochondrial
TGFB: transforming growth factor-beta
TIMP1: tissue inhibitor of matrix metalloprotease-1
TNF-α: tumor necrosis factor alpha
WB: western blotting
WT: wild type
ZFN: zinc finger nuclease technology
α-SMA: alpha smooth muscle actin

---

**Figure 1- figure supplemental 1.**
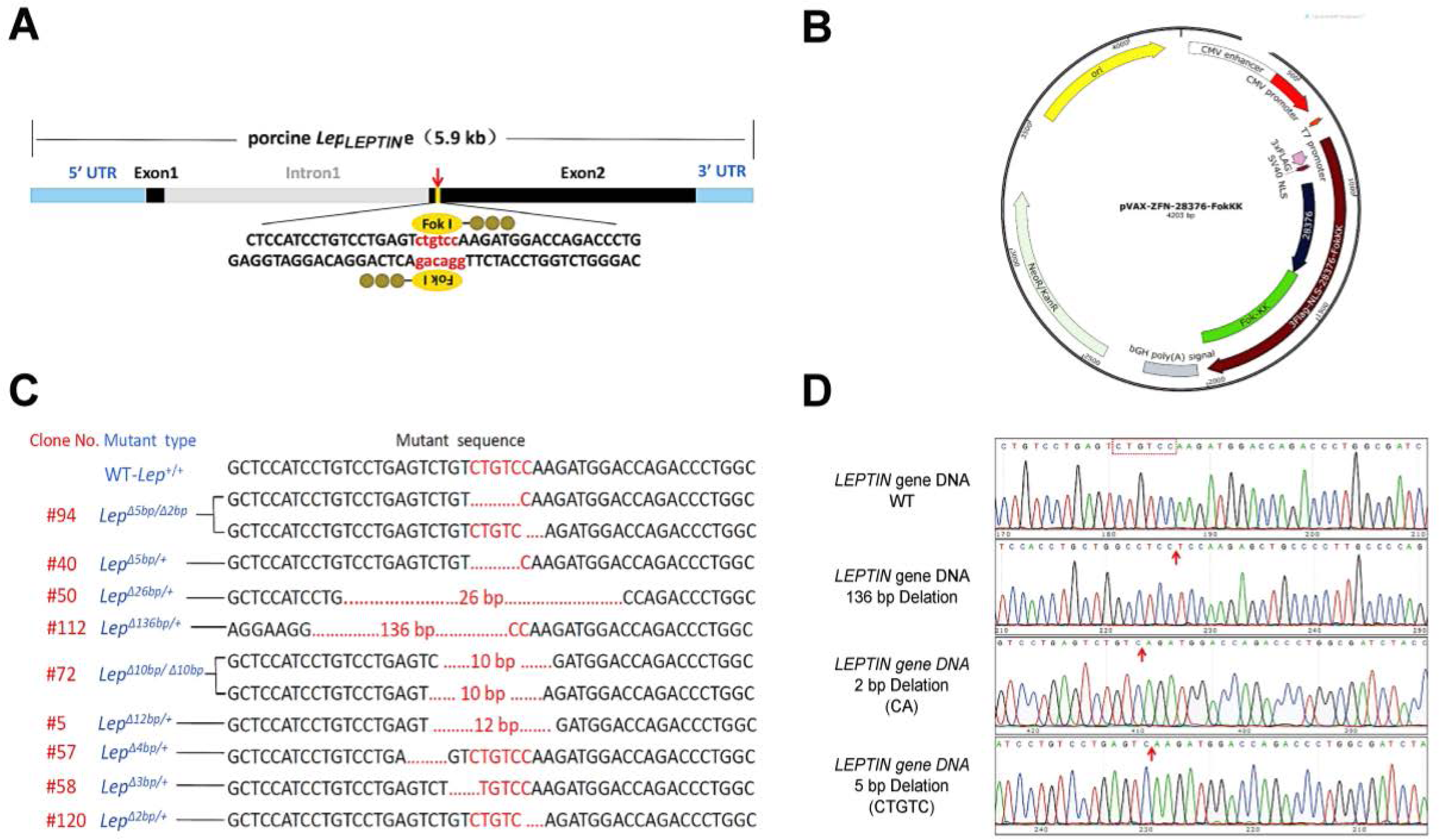
Generation of *LEPTIN* mutant pigs. **A.** Porcine *LEPTIN* gene targeting scheme (targeting site in red). **B.** Construction of ZFN target vector. **C.** Positive clones and observed *LEPTIN* mutations. **D.** Sequence analysis of *LEPTIN* mutants. Peak maps shown mutated base pairs in *LEPTIN* gene on three different chromatids.

**Figure 1- figure supplemental 2.**
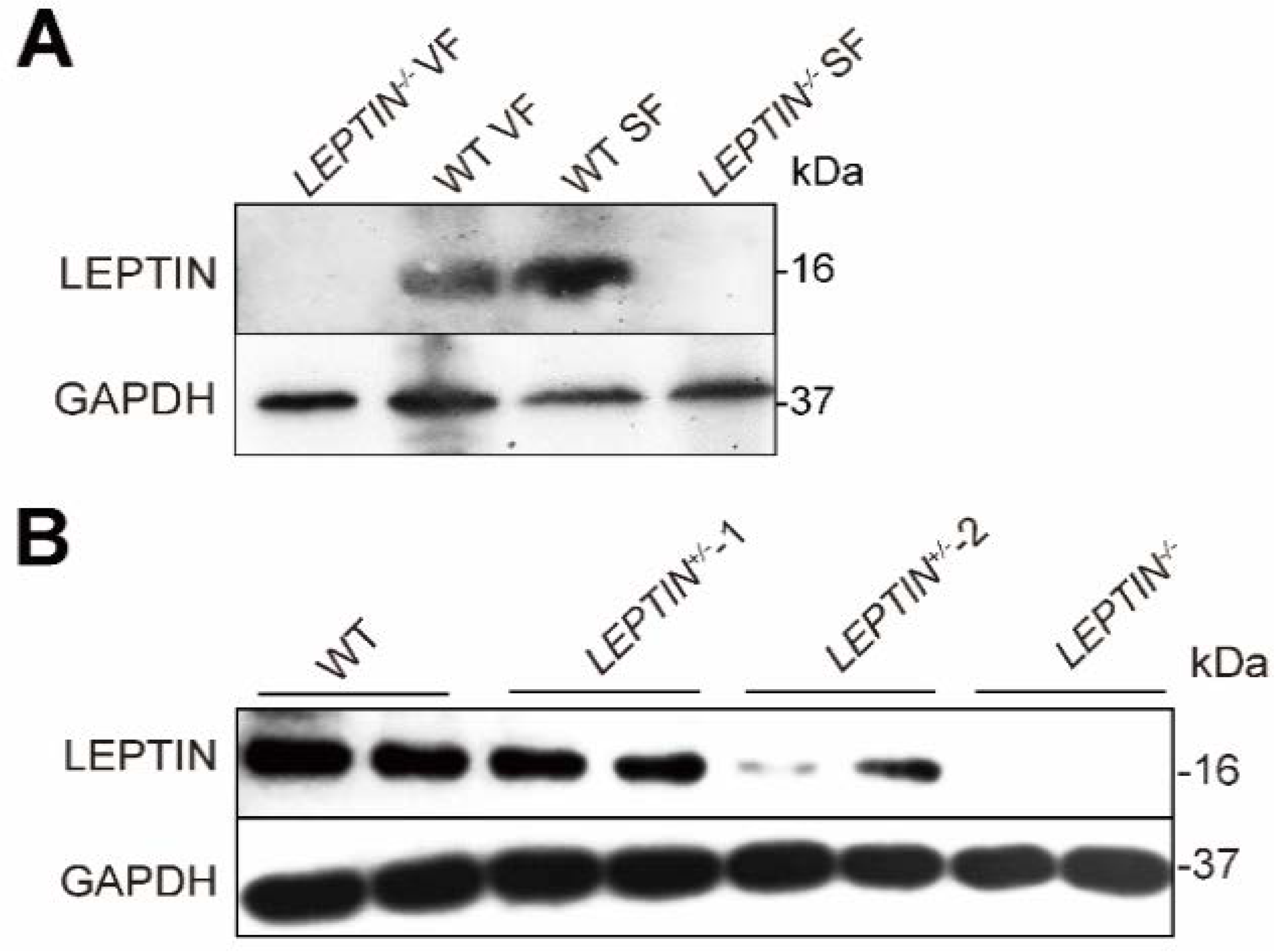
Detection of Leptin in mutant pigs. **A.** The level of Leptin in subcutaneous fat and visceral fat was measured by WB. **B.** WB of Leptin expression in serum.

**Figure 1- figure supplemental 3.**
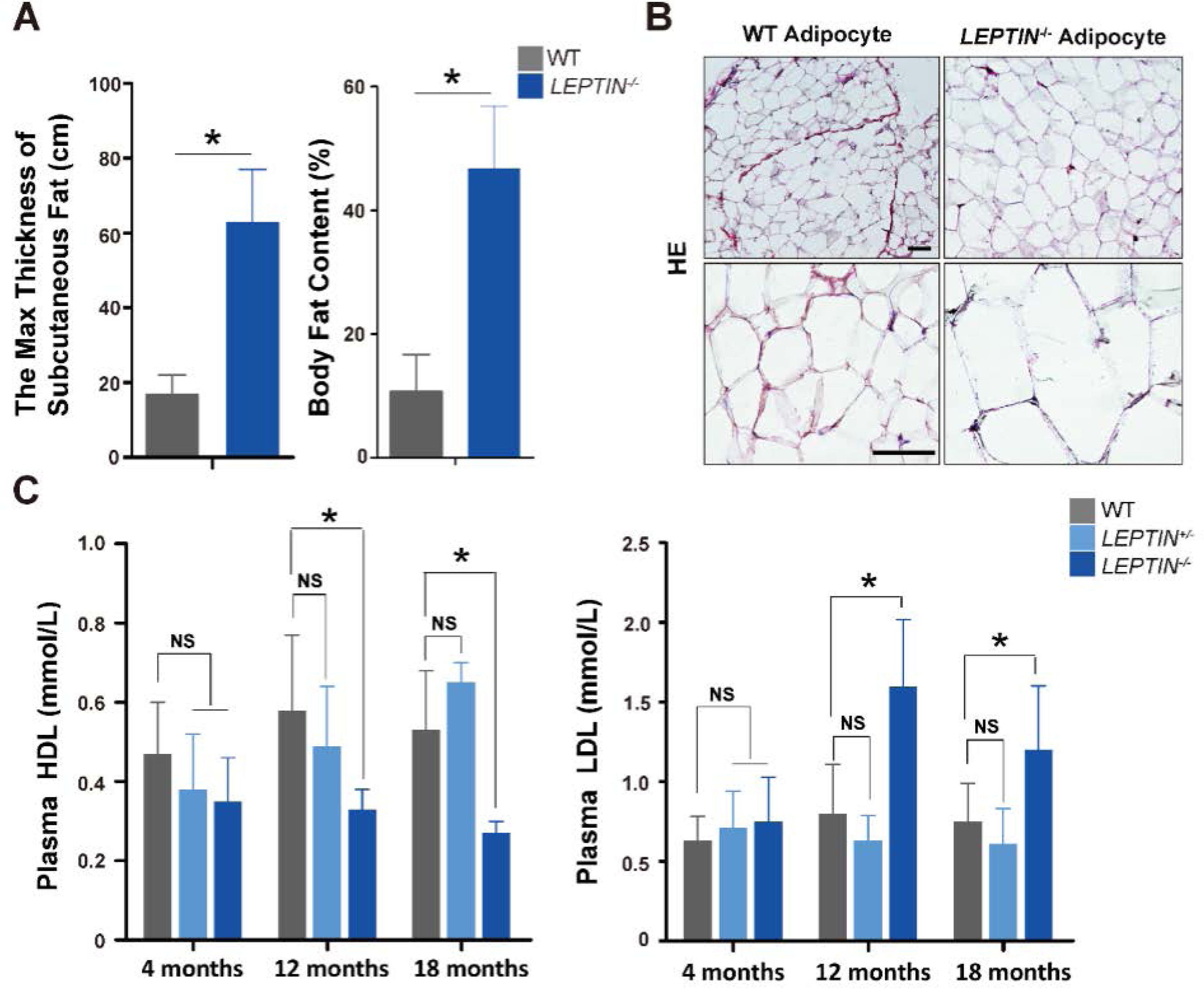
Analysis of obesity phenotypes in *LEPTIN*^-/-^ pigs. **A.** Maximum thickness of subcutaneous fat and the percentage of body fat in *LEPTIN*^-/-^ and WT pigs. **B.** HE staining of adipose tissue. Bar=100μm. **C.** Serum high density lipoprotein (HDL) and low density lipoprotein (LDL) concentrations. The bars represent the mean ± SD; NS, non-significant.*P<0.05. Bar=20μm.

**Figure 1- figure supplemental 4.**
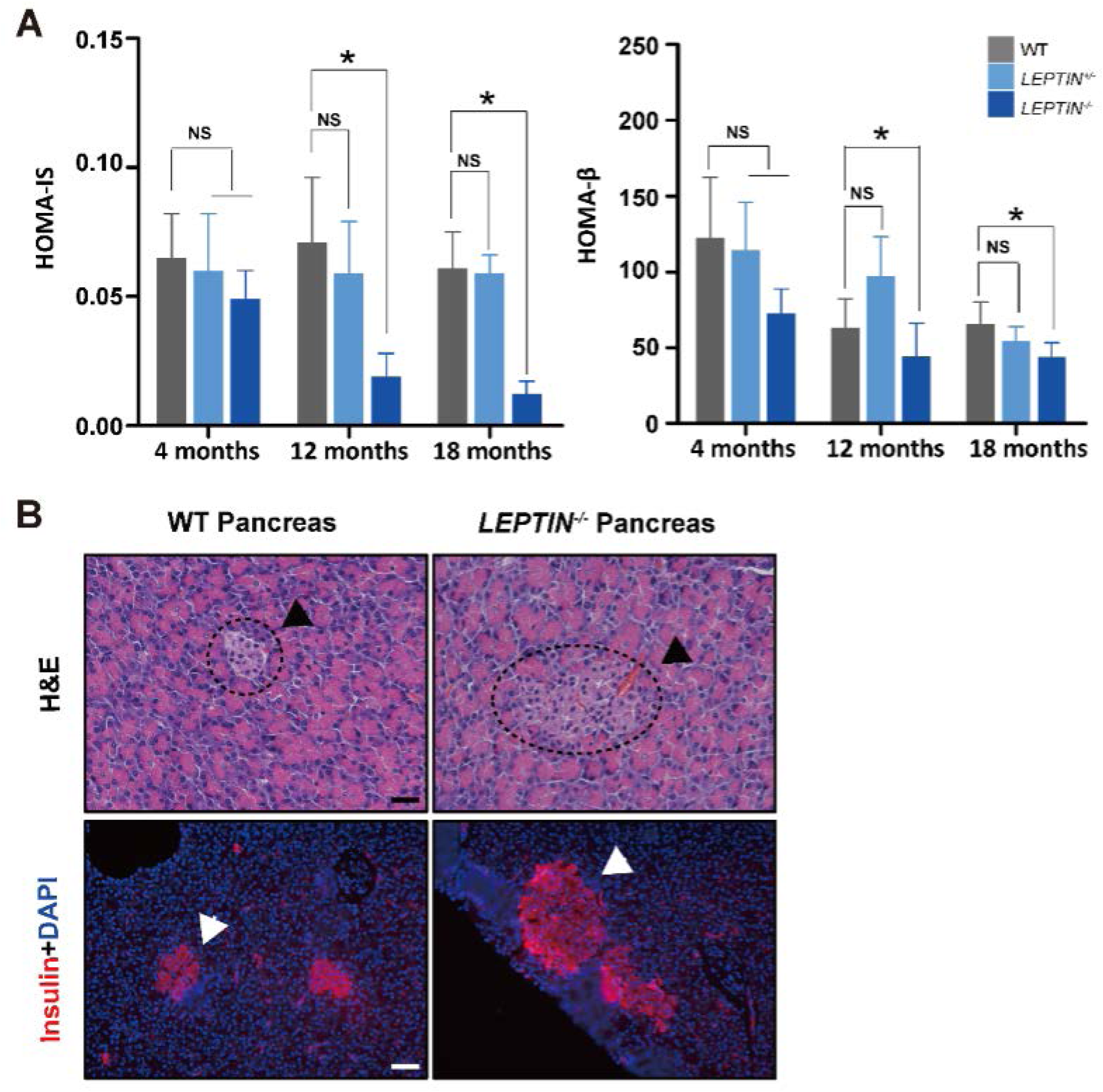
Analysis of the type II diabetes in *LEPTIN*^-/-^ pigs. **A.** HOMA-IS analysis to evaluate insulin sensitivity and HOMA-β analysis to evaluate function of β cells for pigs at different ages. **B.** H&E staining and insulin immuno-fluorescence of pig pancreatic tissue. Dotted box and arrows indicate islet and β cells (insulin positive). The bars represent the mean ± SD; NS, non-significant.*P<0.05. Bar=20μm.

**Figure 4- figure supplemental 1.**
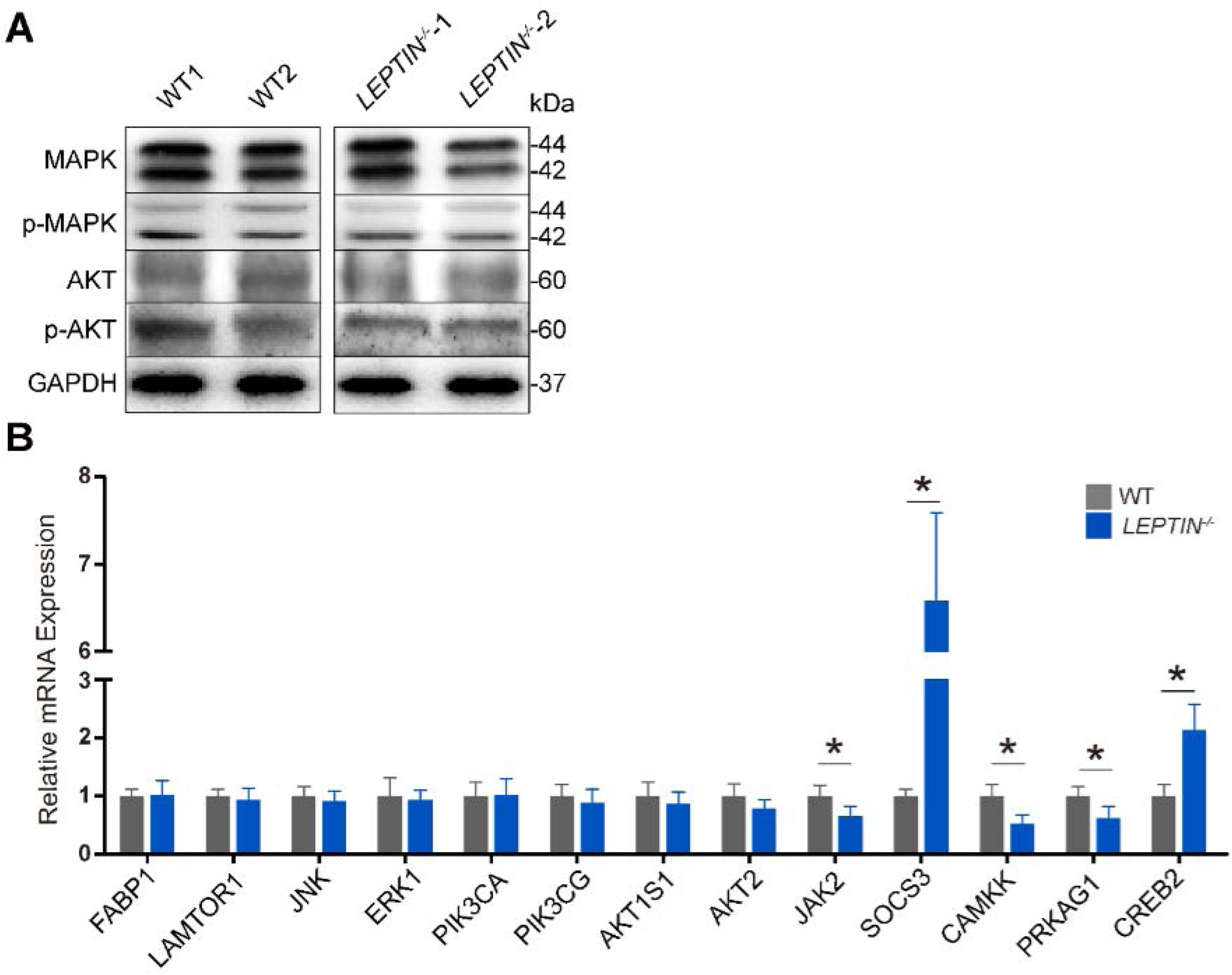
The effects of LEPTIN deficiency on mTOR, MAPK and PI3K-AKT pathways in pig livers. **A.** WB analysis of mTOR, PI3K-AKT, MAPK pathway related proteins**. B.** Expression of mTOR, MAPK, and PI3K-AKT pathway related genes detected by qPCR. The bars represent the mean ± SD; *P<0.05.

**Figure 4- figure supplemental 2.**
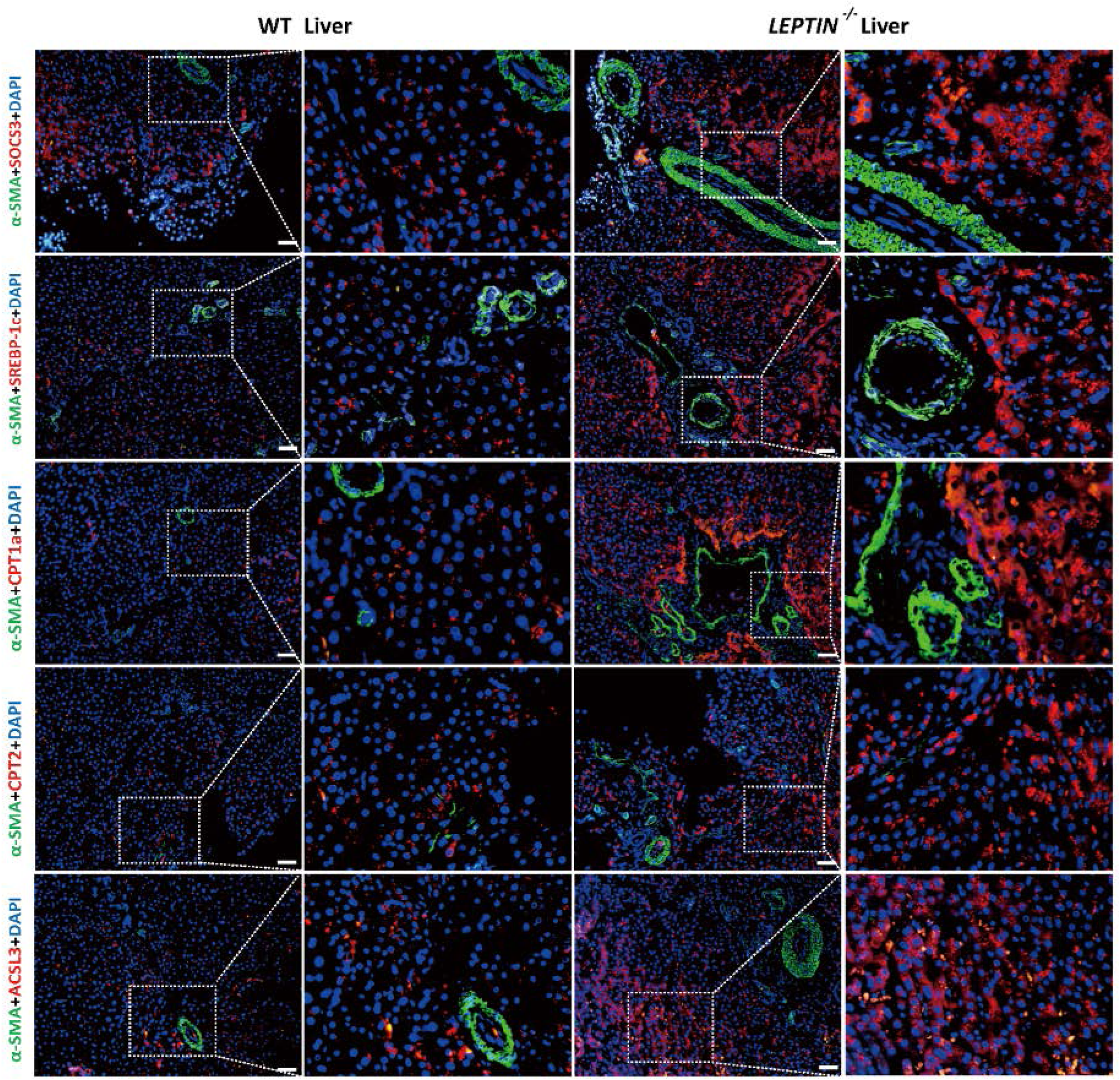
Histological detection of proteins affected by JAK-STAT signaling. Immunofluorescence staining of JAK-STAT pathway related proteins involved in FFA synthesis and β-oxidation process. The white dashed box indicated the enlarged area. Bar=100μm.

**Figure 4- figure supplemental 3.**
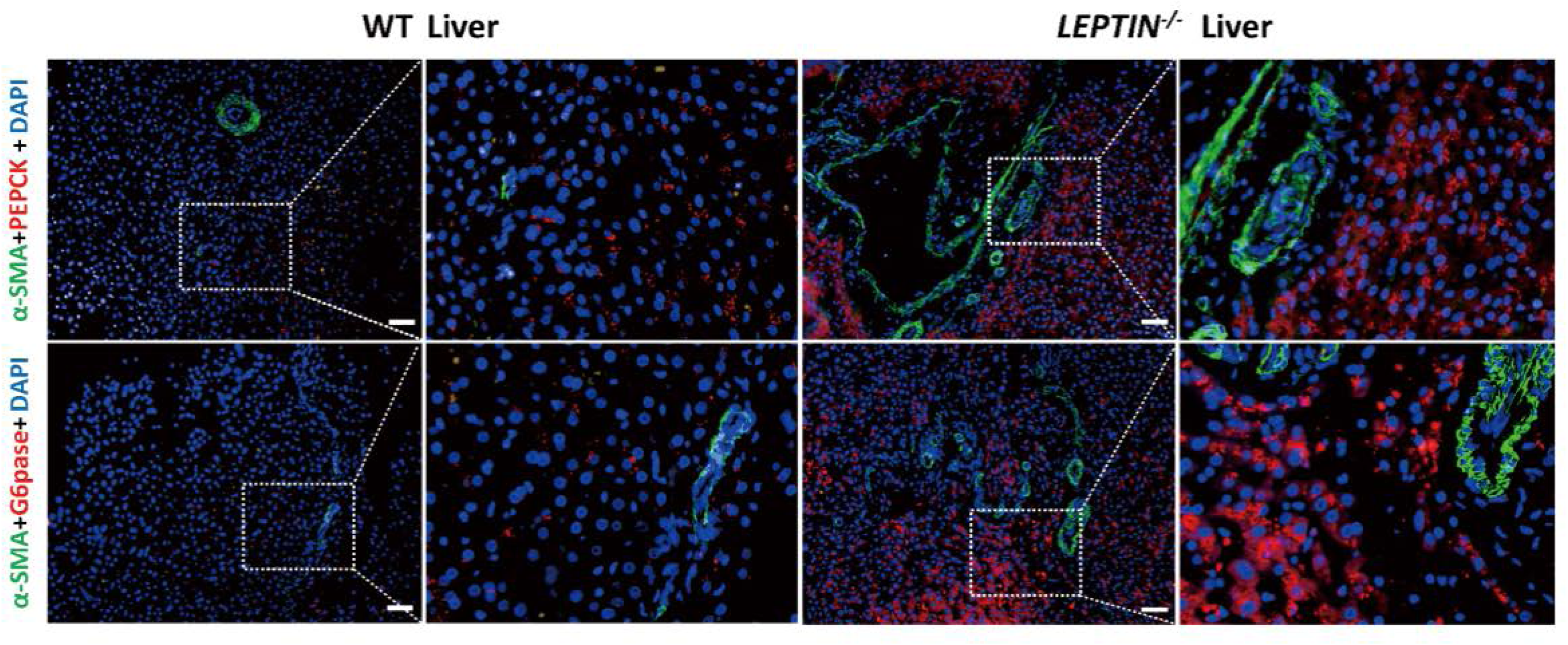
Histological detection of proteins affected by AMPK signaling. Immunofluorescence staining of AMPK pathway related proteins involved in gluconeogenesis. The white dashed box indicated the enlarged area. Bar=100μm.

**Figure 4- figure supplemental 4.**
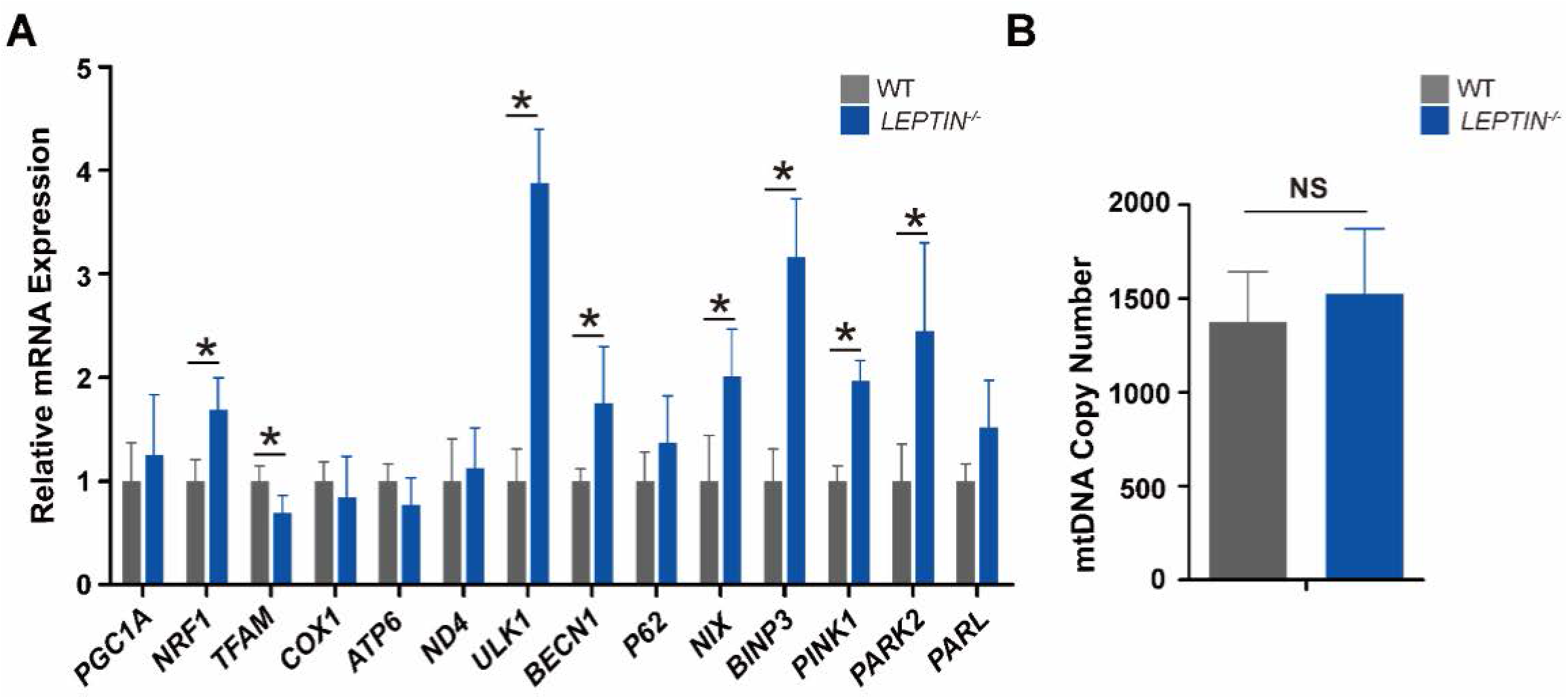
Effects of *LEPTIN* knockout on mitochondrial function in pig livers. **A.** Expression analysis of mitochondrial synthesis and autophagy related genes by qPCR. **B.** Measurement of Mitochondrial DNA copy numbers. The bars represent the mean ± SD; NS, non-significant.*P<0.05.

**Figure 4- source data. Original files of western blot.**

**Figure 5- figure supplemental 1.**
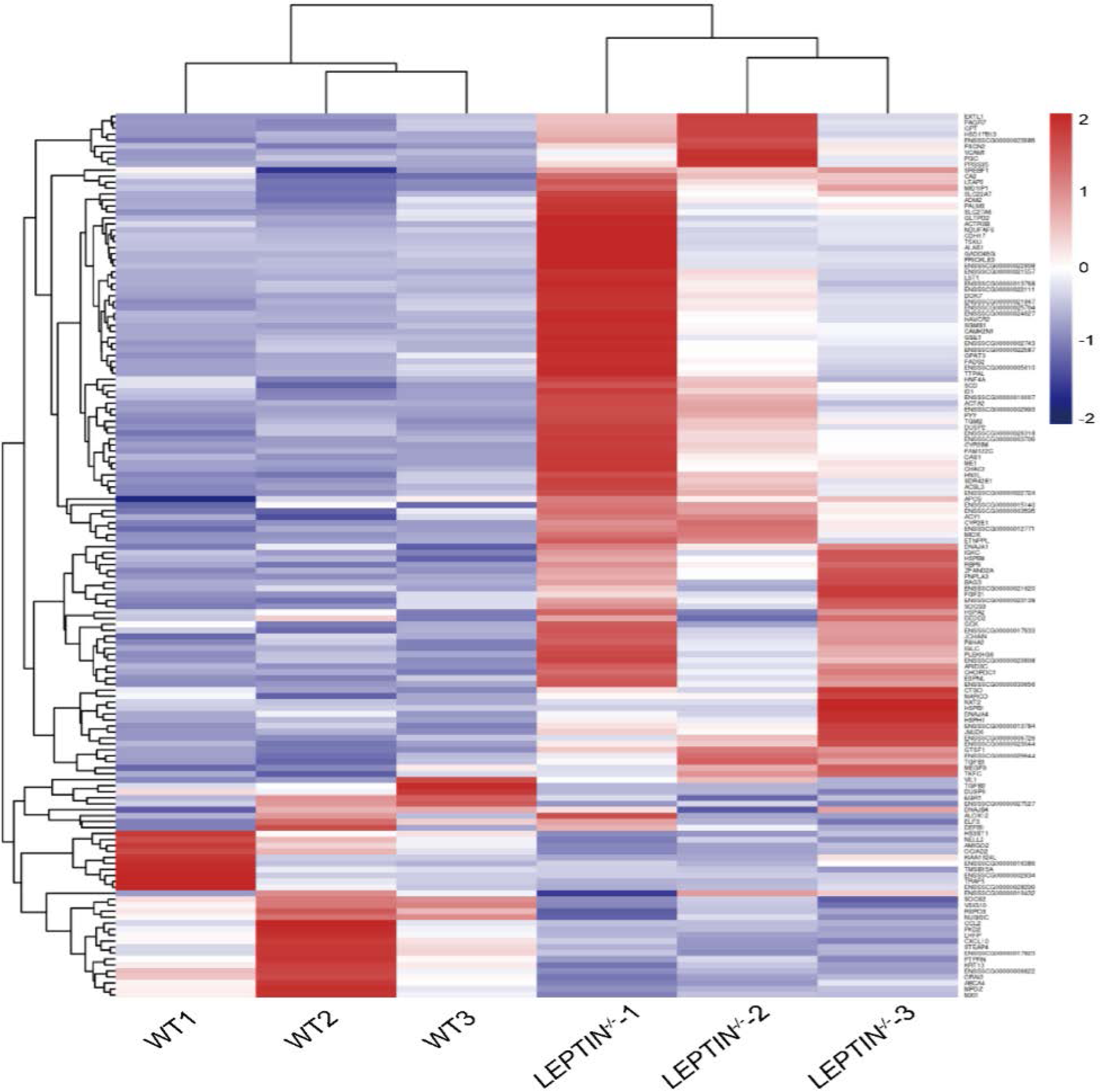
Heatmap of DEGs between *LEPTIN*^-/-^ and WT pig livers. Heatmap displaying the relative expression of DEGs in liver samples from three WT and *LEPTIN*^-/-^ pigs assessed via RNA-seq. Red indicates up-regulated and blue indicates down-regulated genes.

**Figure 5- figure supplemental 2.**
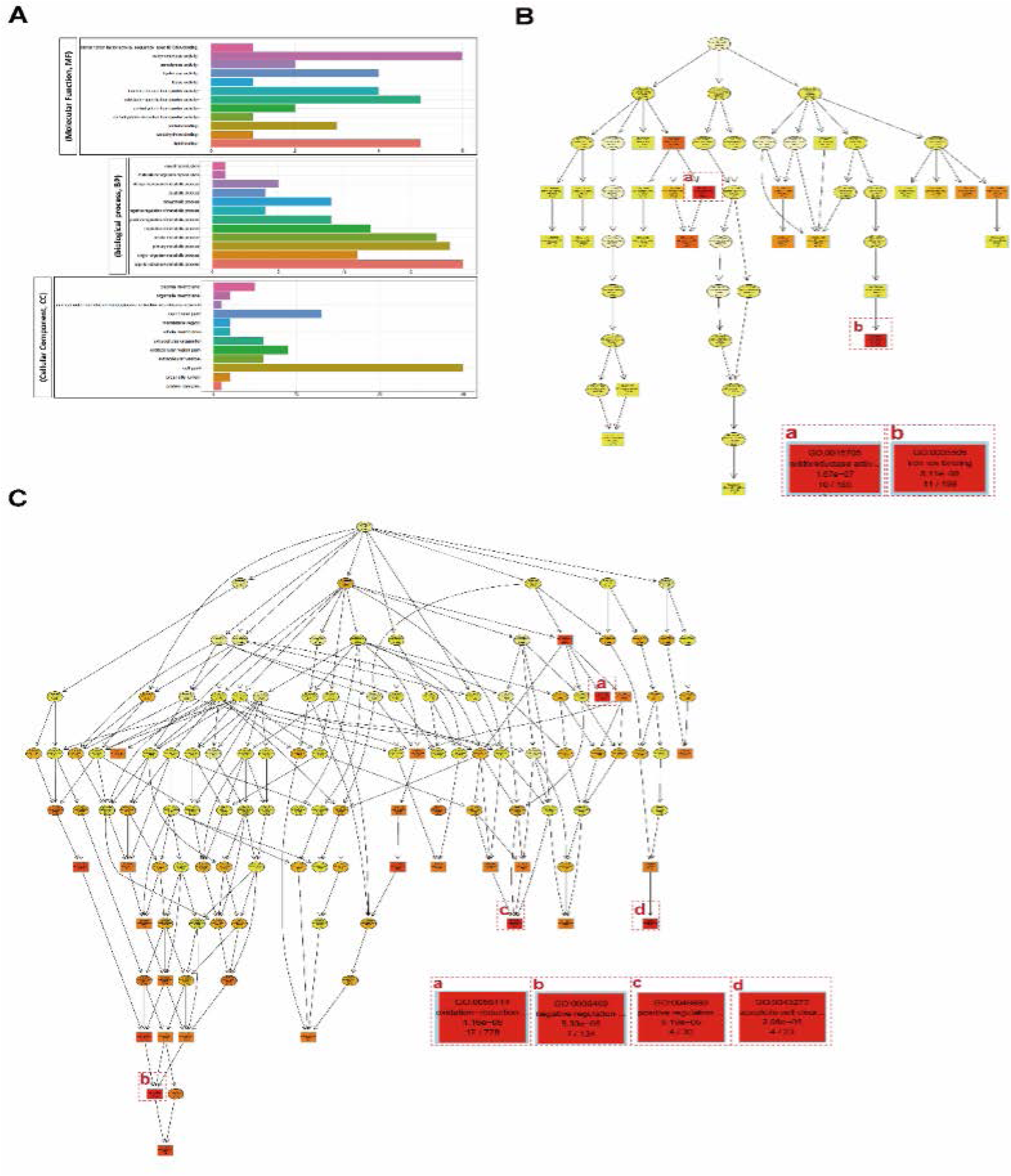
GO terms enriched for DEGs. **A.** Enrichment of molecular function, biological process and cell component GO terms. The ordinate represents the entry names for different functions, processes and components, and the abscissa represents the number of genes enriched on that entry. **B.** Molecular functional pattern of enriched GO terms. Each node represents a GO term, and the darkness of the color indicates the degree of enrichment. The term name and corrected *P*-value are shown for each node. The node terms **(a)** and **(b)** represent the two most highly enriched of the molecular functions. **C.** Biological process pattern of enriched GO terms. The node terms **a**, **b**, **c** and **d** are the four most highly enriched biological processes.

**Figure 5- figure supplemental 3.**
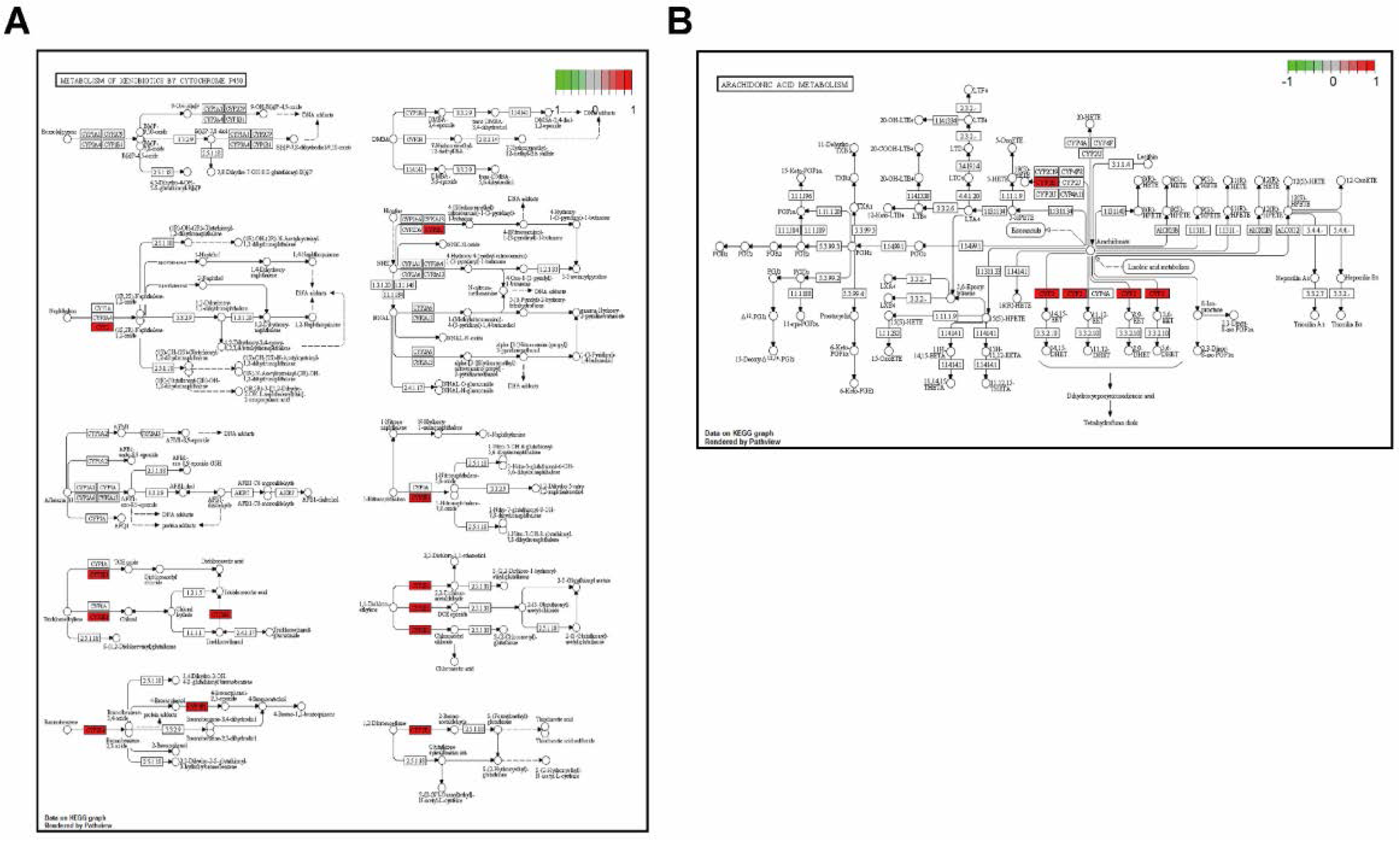
P450 enzyme (CYP2) related KEGG pathways enrichment for DEGs. Boxes represent molecular compounds, solid arrows represent chemical reactions, and dotted arrows represent indirect reactions. Red boxes represent DEGs. **A.** Metabolism of xenobiotics by cytochrome P450 (00980). **B.** Arachidonic acid metabolism pathway (00590).

**Figure 5- source data. Original files of western blot.**

**Figure 6- figure supplemental 1.**
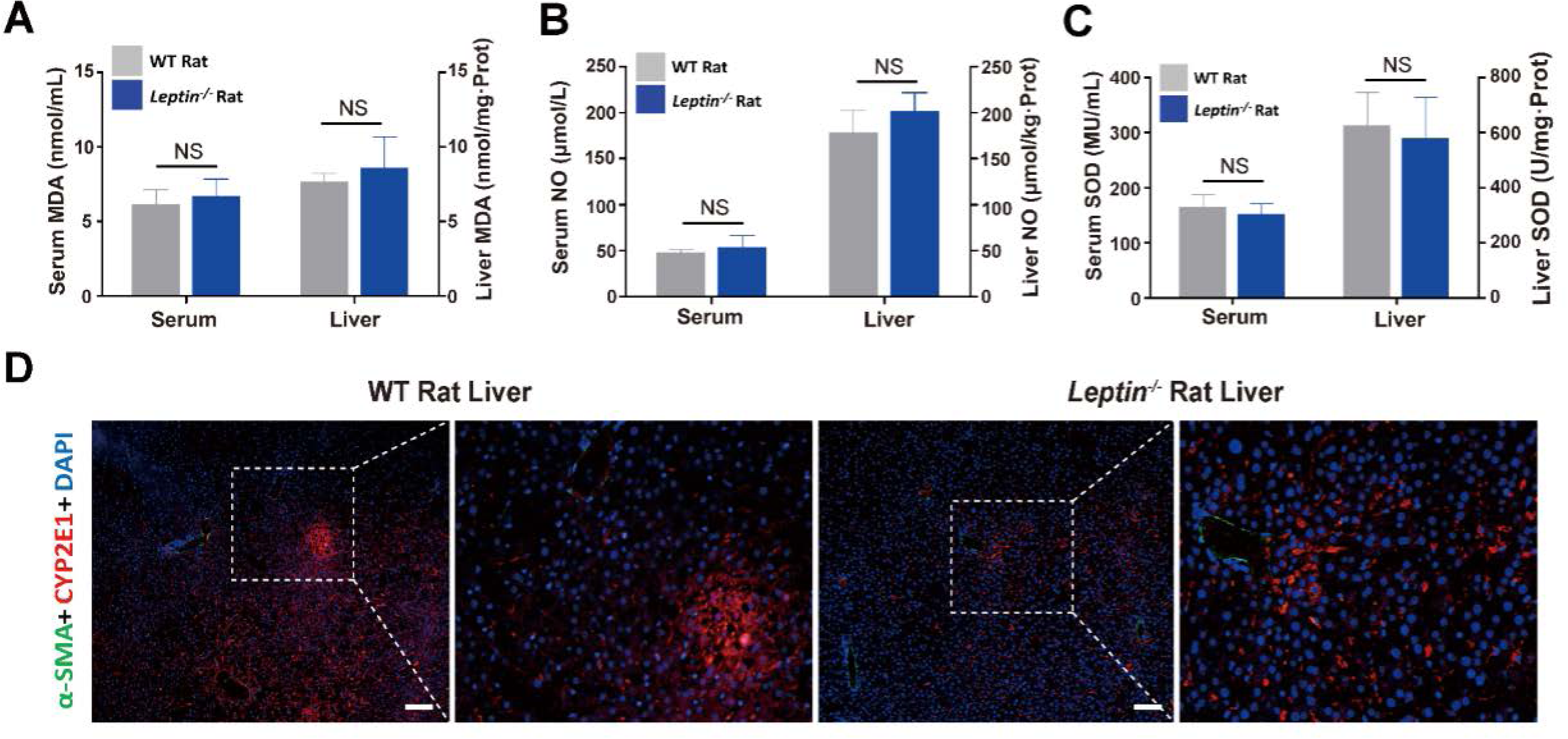
Oxidative stress analysis in *Leptin^-/-^* rat liver. **A-C.** Analysis of oxidative stress markers, MDA **(A),** NO **(B) and** SOD **(C)**, in the *Leptin*^-/-^ rat liver and serum. n=3. The bars represent the mean±SD; NS, non-significant. **D**. Immunofluorescence staining of CYP2E1. The white dashed box indicated the enlarged area. Bar=100μm.

**Figure 6- figure supplemental 2.**
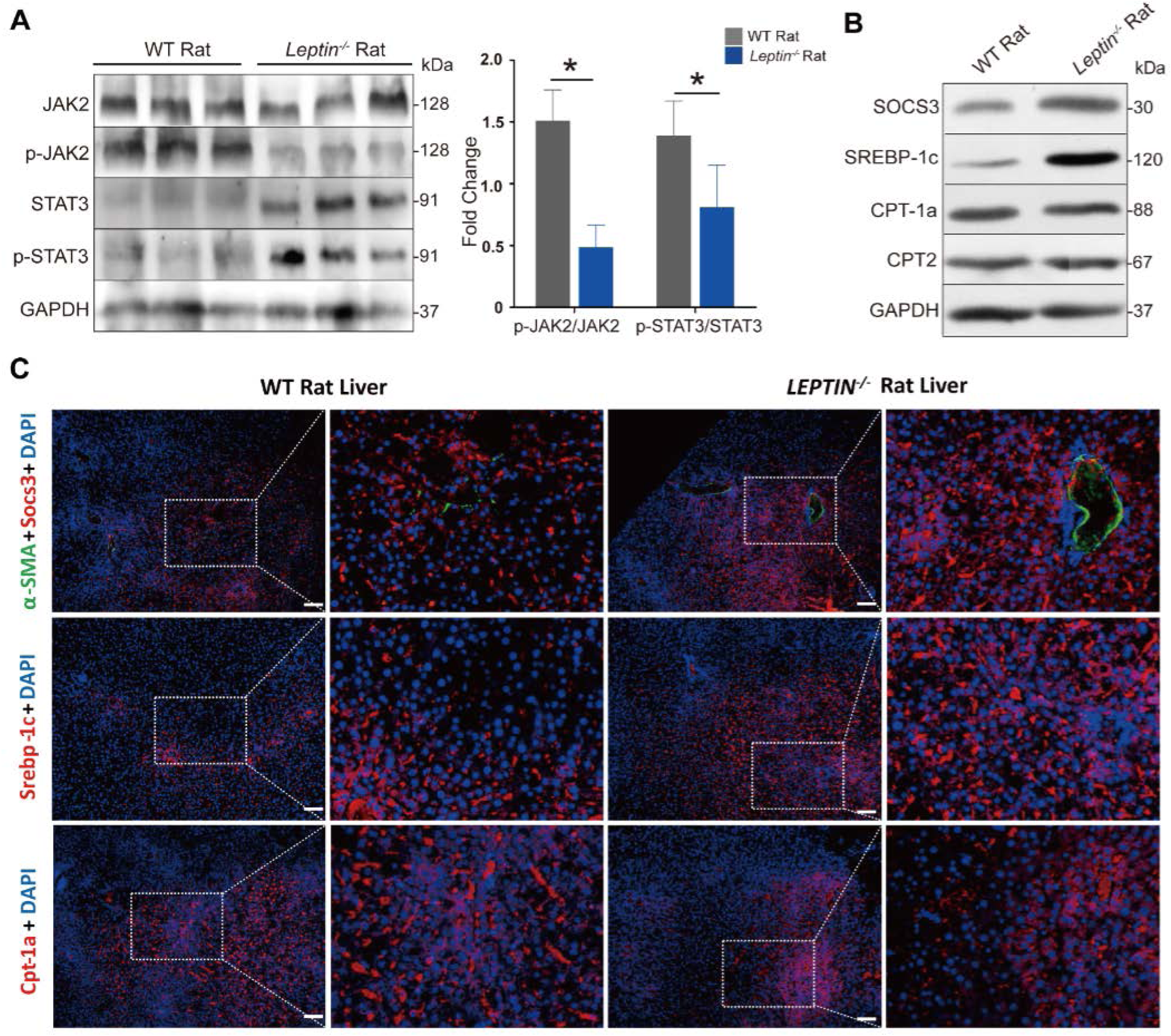
Analysis JAK-STAT pathway related signaling and histological detection of proteins affected by JAK-STAT pathway in *Leptin*^-/-^ rats. **A.** WB analysis of CYP2E1 in rat livers. **B.** WB analysis of JAK-STAT pathway related proteins in rat livers. Gray scale quantitative analysis of JAK2/p-JAK2 and STAT3/p-STAT3 protein expression. The bars represent the mean ± SD; *P<0.05. NS, non-significant. **C.** Immunofluorescence staining of JAK-STAT pathway related proteins involved in FFA synthesis and β-oxidation processes. The white dashed box indicated the enlarged area. Bar=100μm.

**Figure 6- figure supplemental 3.**
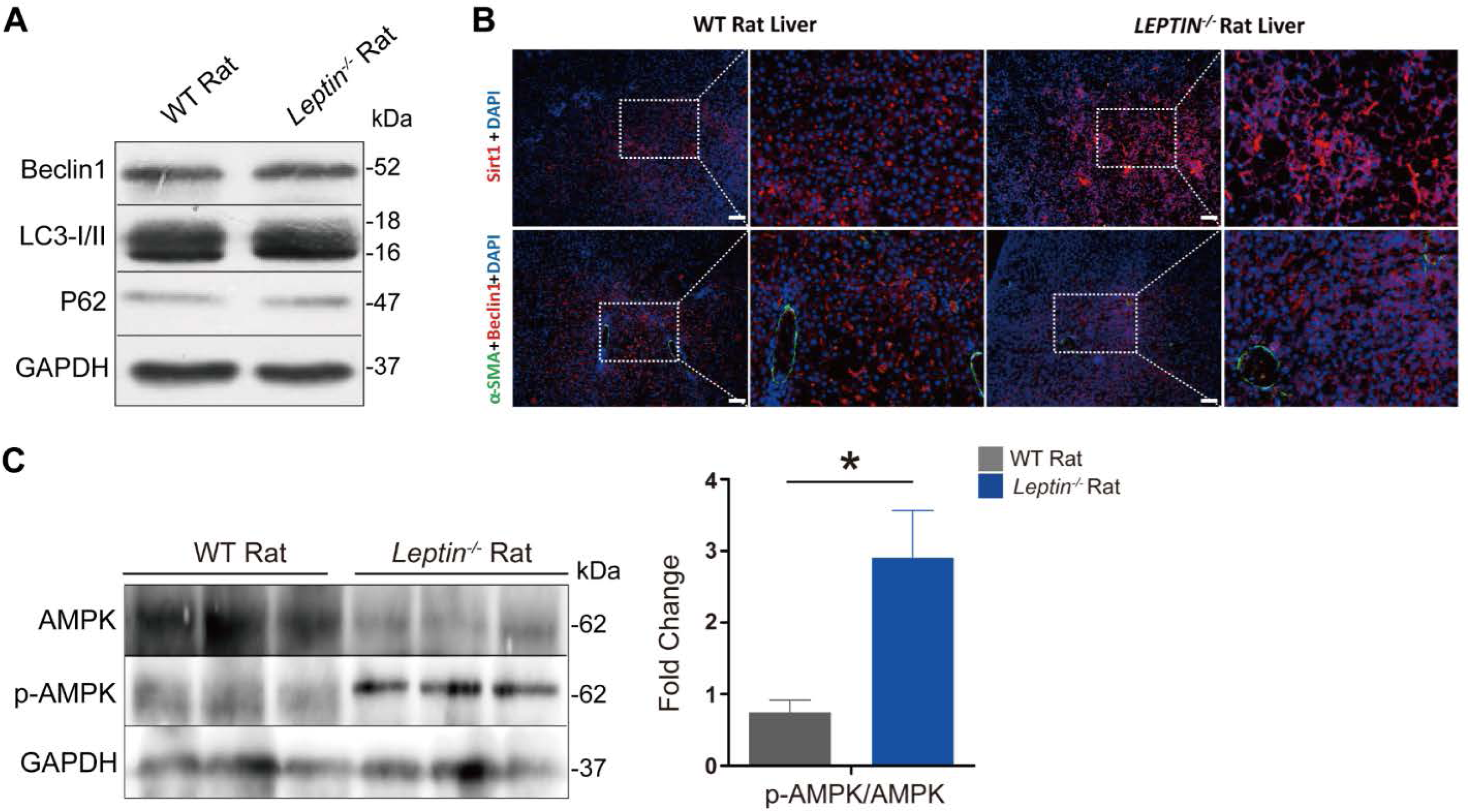
Analysis AMPK pathway related signaling and histological detection of proteins affected by AMPK pathway in *Leptin*^-/-^ rats. **A.** WB analysis of mitochondrial autophagy related markers. **B.** Immunofluorescence staining of AMPK pathway related proteins involved in mitochondrial autophagy. The white dashed box indicated the enlarged area. Bar=100μm. **C.** WB analysis of AMPK and p-AMPK proteins. Gray scale quantitative analysis of AMPK/p-AMPK protein expression. n=3/group. The bars represent the mean±SD; *P<0.05. NS, non-significant.

**Figure 6- source data. Original files of western blot.**

**Supplemental Table 1. Analysis of off-target mutations in *LEPTIN*^-/-^ pigs.**

Sequence locations were obtained by whole pig genome BLAST with ZFN cut site sequences. The yellow part represents the target site section, with each homologous section and the chromosome position on the genome listed below. “No” meant that there was no genetic mutation detected.

**Supplemental Table 2. KEGG pathways were enriched for DEGs.**

The analysis of pathway enrichment was based on the KEGG pathway analysis (*P*< 0.05), and hypergeometric tests were used to screen the pathways with significant enrichment.

**Supplemental Table 3. The list of antibodies used in this study.**

In this table, the item No. and manufacturer of all antibodies used in this study for WB and immunofluorescence staining are listed in details.

**Supplemental Table 4. The list of primers used in this study.**

In this table, the sequence of all primers used in the study for quantitative PCR are listed in details.

